# Spatial structure impacts adaptive therapy by shaping intra-tumoral competition

**DOI:** 10.1101/2020.11.03.365163

**Authors:** Maximilian A. R. Strobl, Jill Gallaher, Jeffrey West, Mark Robertson-Tessi, Philip K. Maini, Alexander R. A. Anderson

**Author notes:** These authors jointly supervised this work (Philip K. Maini, Alexander R. A. Anderson).

## Abstract

(1) Background: Adaptive therapy aims to tackle cancer drug resistance by leveraging intra-tumoral competition between drug-sensitive and resistant cells. Motivated by promising results in prostate cancer there is growing interest in extending this approach to other cancers. Here we present a theoretical study of intra-tumoral competition during adaptive therapy, to identify under which circumstances it will be superior to aggressive treatment; (2) Methods: We use a 2-D, on-lattice, agent-based tumour model to examine the impact of different microenvironmental factors on the comparison between continuous drug administration and adaptive therapy. (3) Results: We show that the degree of crowding, the initial resistance fraction, the presence of resistance costs, and the rate of tumour cell turnover are key determinants of the benefit of adaptive therapy, and we study in detail how these factors alter competition between cells. We find that intra-specific competition between resistant cells plays an unexpectedly important role in the ability to control resistance. To conclude we show how differences in resistance cost and turnover change the tumour’s spatial organisation and may explain differences in cycling speed observed in a cohort of 67 prostate cancer patients undergoing intermittent androgen deprivation therapy; (4) Conclusion: Our work provides insights into how adaptive therapy leverages inter- and intra-specific competition to control resistance, and shows that the tumour’s spatial architecture will likely be an important factor in determining the quantitative benefit of adaptive therapy in patients.

## 1. Introduction

“Can insects become resistant to sprays?” - this is the question entomologist Axel Melander raised in an article of the same title in 1914 [1]. At a site in Clarkston, WA, Melander had observed that over 90% of an insect pest called the “San Jose scale” was surviving despite being sprayed with sulphur-lime insecticide [1]. If we were to ask the same question in cancer treatment today we would be met with an equally resounding “yes”: while for most cancers it is possible to achieve an initial, possibly significant, burden reduction, many patients recur with drug-resistant disease, or even progress while still under treatment. Drug resistance can develop in a number of ways, including genetic mutations which alter drug binding, changes in gene expression which activate alternative signalling pathways, or environmentally-mediated resistance [2-4]. In the clinic, the main strategy for managing cancer drug resistance is to switch treatment with the aim of finding an agent to which the tumour is still susceptible [3,4]. Similarly, Melander suggested it might be possible to tackle sulphur-lime resistance by switching to oil-based sprays [1]. But he also foresaw the possibility of, and the challenges arising from, multi-drug resistance [1]. As an alternative, Melander proposed that it might be possible to maintain insecticide sensitivity through less aggressive spraying as this would promote inter-breeding of sensitive and resistant populations and, thus, dilute the resistant genotype [1]. Allocation of drug-free “refuge” patches in the neighbourhood of plots in which an insecticide is used is one modality of modern pest management and is even required by law for the use of certain agents in the US (e.g. Bt-crops [5]).

Recently, the concept that treatment de-escalation can delay the emergence of resistance has found application also in oncology. Standard-of-care cancer treatment regimens aim to maximise cell kill through application of the maximum tolerated dose (MTD), in order to achieve a cure. In contrast, an emerging approach called adaptive therapy proposes to focus not on burden reduction, but on burden control in settings, such as advanced, metastatic disease, in which cures are unlikely [6—9]. Eradication strategies free surviving cells from intra-tumoral resource competition which would otherwise inhibit resistance growth. Adaptive therapy aims to leverage this competition by maintaining drug-sensitive cells in order to avoid, or at least delay, the emergence of resistance [6,8]. A number of pre-clinical studies have demonstrated the feasibility of this approach in ovarian [7], breast [10], colorectal [11], and skin cancer [12]. Moreover, a clinical trial of adaptive therapy in metastatic castration resistant prostate cancer achieved not only an at least 10 month increase in median time to progression (TTP), but also a 53% reduction in cumulative drug usage [13]. Further clinical trials in castration sensitive prostate cancer and melanoma are ongoing (clinicaltrials.gov identifiers NCT03511196 and NCT03543969, respectively).

In addition to testing its feasibility, there has been significant interest in characterising the underpinning eco-evolutionary principles of adaptive therapy through mathematical modelling. We identify three key results. The first insight was derived from approaches which represent the tumour as a mixture of drug-sensitive and resistant cells modelled as a system of two or more ordinary differential equations (ODEs) with competition described by the Lotka-Volterra model from ecology [7,11,12,14–17] or by a matrix game [18]. These analyses have demonstrated that less aggressive treatment allows for longer tumour control under a range of assumptions on the tumour growth law (exponential: [11,14,16,18]; logistic: [7,12,14,16]; Gompertzian: [14–16]; dynamic carrying capacity: [11,17]), and the origin of resistance (pre-existing: [7,11,14-16,18]; acquired [12,15,16]; cancer stem-cell-based: [17]). Furthermore, this work predicts that adaptive therapy will be most effective in cases where cures are unlikely due to pre-existing resistance and where at the same time conditions (resistance fraction, proximity to carrying capacity) are such that inter-specific competition with drug-sensitive cells is strong (see [16] for a comprehensive and formal summary of these results). The second key result is that while these conclusions broadly transfer to more complex, spatially-explicit tumour models, the spatial distribution of resistant cells is important [11,19]. Bacevic et al [11] showed in a two-dimensional (2-D), on-lattice, agent-based model (ABM) of a tumour spheroid that longer control is achieved if resistance arises in the centre of the tumour compared to when it arises on the edge. Gallaher et al [19] corroborated this result in a 2-D, off-lattice setting with resistance modelled as a continuum, and further demonstrated that tumour control was adversely affected by high cell motility and cell plasticity. Thirdly, models focussed on metastatic prostate cancer have illustrated how these concepts may be realised in a specific disease pathology [13,17] and how we may enhance tumour control by using a multi-drug approach [17,20,21].

Using a minimalistic, sensitive/resistant Lotka-Volterra ODE model we recently examined the role of two factors not previously considered: resistance costs and cellular turnover [22]. The “cost of resistance” refers to the notion that drug-resistance may come at a fitness cost, such as a decreased proliferation rate, in a drug-free environment due to, for example, increased energy consumption. Such costs have been reported amongst others in colorectal cancer [11], breast cancer [19], and melanoma [12], and have been assumed to be necessary for the success of adaptive therapy [6,8]. We showed that this is not the case, and furthermore that the impact of a resistance cost depends on the rate of tumour cell death. Moreover, we found that higher rates of tumour cell death increased the benefit of adaptive therapy more generally as they amplified the effects of competition: the shorter a cell’s lifespan and the fewer opportunities for division, the greater the impact if proliferation is inhibited by competition [22].

The goal of this paper is to examine whether these conclusions still hold true if space is taken into account. In addition, we aim to identify a “minimal spatial working model” for adaptive therapy, as both previous spatial models have made assumptions about nutrient supply [11] or cell cycle dynamics [19] which while making these models more realistic, does complicate identification of governing principles. In the following, we recast our previously derived ODE model into a 2-D, on-lattice, ABM in which the tumour is assumed to be composed of drug-sensitive and resistant cells. We examine the roles of the initial resistance fraction, the proximity of the tumour to carrying capacity, resistance costs and cellular turnover and confirm that the behaviour we observe is consistent with our prior non-spatial results. Furthermore, our analysis shows that the spatial distribution of resistant cells is a key factor in determining a tumour competition profile and modulates TTP, the benefit of adaptive therapy, and the variability in outcomes. In particular, we identify intra-specific competition between resistant cells at the edge and core of a growing colony as an important factor in adaptive therapy. We conclude by applying our insights to intermittent androgen deprivation therapy in prostate cancer, where we show that distinct underlying spatial architectures, driven by differences in resistance costs and turnover, may explain observed differences in cycling speed between 67 patients in a Phase II trial by Bruchovsky et al [23]. Overall, our work helps to provide a more detailed understanding of the spatial competition between sensitive and resistant cells during adaptive therapy and shows that the spatial architecture of the tumour can significantly affect treatment outcomes.

## 2. Materials and Methods

### 2.1. The mathematical model

Random geno- and pheno-typic variation produces tumour cells which show a degree of drug-resistance even prior to drug exposure. This may manifest as an increased ability to persist and adapt to adverse conditions such as drug exposure, or, though perhaps more rarely, it may take the form of fully developed resistance [3,4]. Selective expansion and further adaptation of this population is thought to be the cause of treatment failure in patients.

To study the evolutionary dynamics in response to treatment we consider a 2-D, on-lattice, ABM representative of a small region of tumour tissue or a metastatic site. For simplicity we assume that we can divide cells into drug-sensitive or fully drug-resistant subpopulations (Figure 1a). We choose an on-lattice, agent-based representation as it allows us to explore the role of space and cell-scale stochasticity in a tractable, yet generalisable, way. Each cell occupies a single site in an l × l square lattice with no-flux boundary conditions, and behaves according to the following rules (Figure 1):

**Figure 1.**
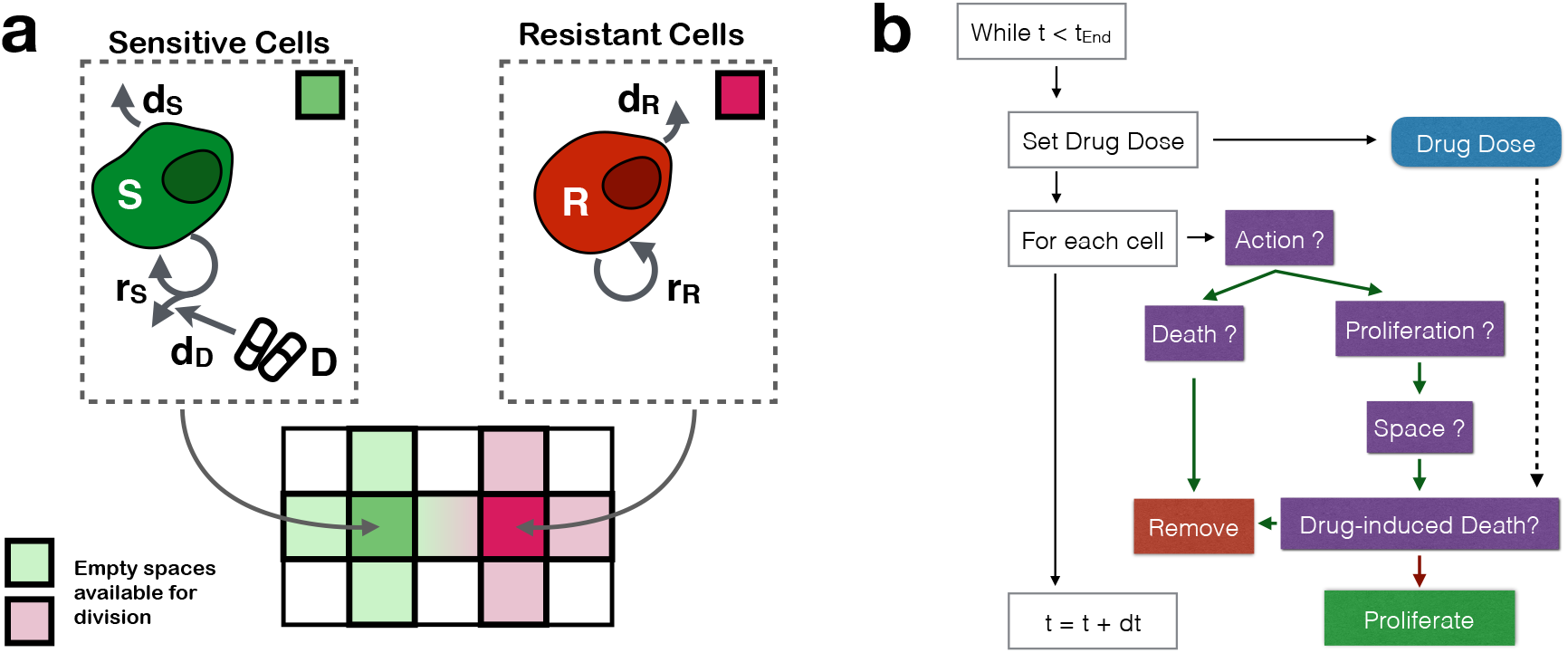
The agent-based tumour model: (**a**) The tumour is modelled as a mixture of drug-sensitive and resistant cells, where each cell occupies a square on a 2-D equi-spaced lattice. Cells divide and die at constant rates. Daughter cells are placed in empty squares in a cell's von Neumann neighbourhood. Drug will kill dividing sensitive cells at a probability *d_D_*. (**b**) Flow diagram of the simulation algorithm.

- Initially, there are a total of *N_0_* cancer cells spread randomly in the tissue, of which a fraction *f_R_* is resistant.
- Sensitive and resistant cells attempt to divide at constant rates *r_s_* and *r_R_* (in units: day^-1^), respectively. If there is at least one empty site in the cell’s von Neumann neighbourhood, then the cell will divide and the daughter will be placed randomly in one of the empty sites in the neighbourhood.
- Cells die at a constant rate *d_T_*. For simplicity, we will assume that both sensitive and resistant cells die at the same rate, *ds = d_R_ = dT*.
- Movement of cells is neglected.
- The domain is sufficiently small so that drug concentration *D(t)* ∈ [0, *D*_Max_] is assumed to be spatially homogeneous throughout the tissue, where *D*_Max_ is the MTD.
- A sensitive cell which is currently undergoing mitosis - that is, it has attempted division and has space available in its neighbourhood - is killed by drug with probability *d_D_D(t)*, where *^d^D* < *D*_Max_.
- Dead cells are immediately removed from the domain.

We denote the number of cells in each population at time *t* by *S(t)* and *R(t),* and the total number by *N(t) = S(t)* + *R(t),* respectively (Table 1).

**Table 1.**
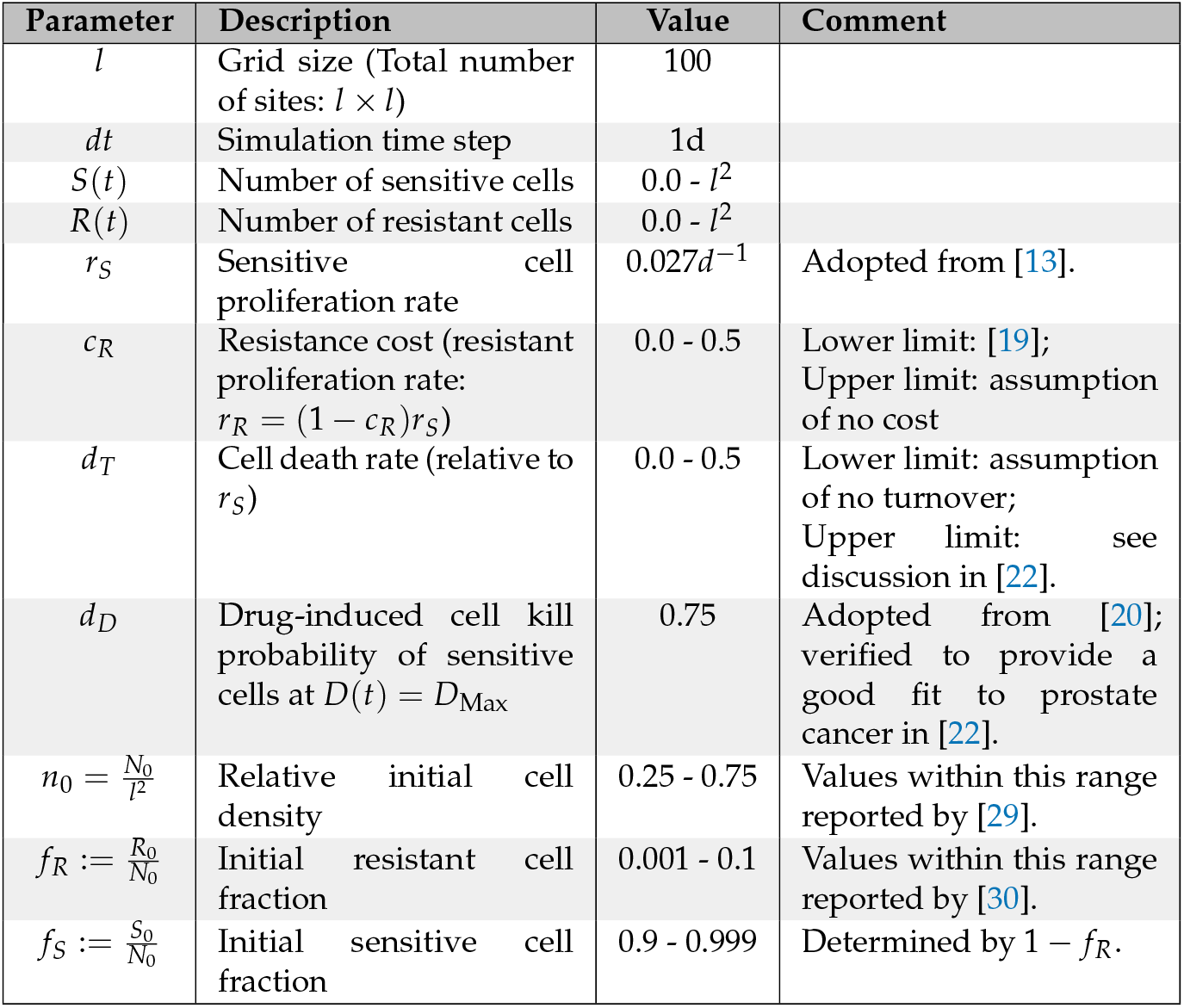
Summary of mathematical model variables and parameters.

| Parameter | Description | Value | Comment |
| --- | --- | --- | --- |
| $l$ | Grid size (Total number of sites: $l \times l$) | 100 | |
| $dt$ | Simulation time step | 1d | |
| $S(t)$ | Number of sensitive cells | $0.0 - l^2$ | |
| $R(t)$ | Number of resistant cells | $0.0 - l^2$ | |
| $r_S$ | Sensitive cell proliferation rate | $0.027d^{-1}$ | Adopted from [13]. |
| $c_R$ | Resistance cost (resistant proliferation rate: $r_R = (1 - c_R)r_S$) | 0.0 - 0.5 | Lower limit: [19]; Upper limit: assumption of no cost |
| $d_T$ | Cell death rate (relative to $r_S$) | 0.0 - 0.5 | Lower limit: assumption of no turnover; Upper limit: see discussion in [22]. |
| $d_D$ | Drug-induced cell kill probability of sensitive cells at $D(t) = D_{\text{Max}}$ | 0.75 | Adopted from [20]; verified to provide a good fit to prostate cancer in [22]. |
| $n_0 = \frac{N_0}{l^2}$ | Relative initial cell density | 0.25 - 0.75 | Values within this range reported by [29]. |
| $f_R := \frac{R_0}{N_0}$ | Initial resistant cell fraction | 0.001 - 0.1 | Values within this range reported by [30]. |
| $f_S := \frac{S_0}{N_0}$ | Initial sensitive cell fraction | 0.9 - 0.999 | Determined by $1 - f_R$. |

We consider two treatment schedules:

1. Continuous therapy at MTD: 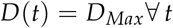.
2. Adaptive therapy as implemented in the Zhang et al [13] prostate cancer clinical trial: Treatment is withdrawn once a 50% decrease from the initial tumour size is achieved, and is reinstated if the original tumour size (*N*_0_) is reached:

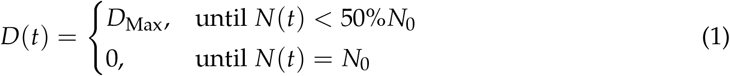

A flow-chart of our model is shown in Figure 1b, and further implementation details are given in Appendix A. We checked convergence (not shown), and performed a consistency analysis [24, 25]. This showed that a sample size upward of 250 provides a representative sample size for our stochastic simulation algorithm (see Appendix B for details). The model is implemented in Java 1.8. using the Hybrid Automata Library [26]. Data analysis was carried out in Python 3.6, using Pandas 0.23.4, Matplotlib 2.2.3, Seaborn 0.9.0, and openCV 3.4.9. The time-evolution of the resistant cells’ neighbourhood composition was visualised using EvoFreq [27] in R 4.0.2. The code will be made available on-line upon publication.

### 2.2. Comparison with the non-spatial model

To understand the impact of space we compare the ABM to the following ODE model which we have studied previously in [22]:

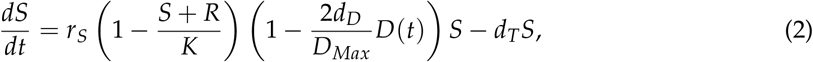

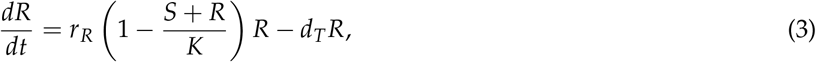

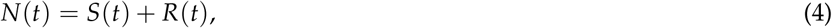

where *K* is the carrying capacity, and the initial conditions are given by *S*(0) = *S*_0_, *R*(0) = *R*_0_, and *N*_0_ = *S*_0_ + *R*_0_, respectively. We set *K = l*^2^ and used the same parameter values as for the agent-based simulation otherwise (Table 1). The equations were solved using the RK5(4) explicit Runge-Kutta scheme [28] provided in Scipy (for further details see [22]).

### 2.3. Model parameters

We parameterise our model using values from the literature for prostate cancer (Table 1). We want to stress, however, that the aim of our work is to develop qualitative understanding, not to make quantitative predictions directed at prostate cancer. As such, our predictions should be interpreted not in an absolute (“treatment X will achieve a TTP of 8m”), but in a relative fashion (“treatment X will achieve a longer TTP than treatment Y because of mechanism Z”).

### 2.4. Analysis of patient data

In order to examine whether our model can explain differences in the cycling speed of patients undergoing intermittent androgen deprivation therapy, we fit it to the publicly available, longitudinal response data from the Phase II trial by Bruchovsky et al [23]. The data were downloaded from http://www.nicholasbruchovsky.com/clinicalResearch.html in July 2020. So as to avoid potentially confounding effects from a change in the number of lesions, patients who developed a metastasis were excluded from analysis. This yielded data from a total of 67 patients. The model was fitted to the normalised PSA measurements by minimising the root mean squared error between the data and the predictions. Given the stochastic nature of the ABM, each candidate fit was assembled from 25 independent stochastic replicates. Optimisation was carried out using the basin-hopping algorithm in Scipy [31] using default search parameters and a maximum of either 50 (2 parameter models) or 75 optimisation steps (4 parameter model). To escape potential local minima, optimisation was repeated 10 times for each patient from different, randomly chosen initial conditions, with only the best fit according to the Akaike Information Criterion taken forward for analysis. We fitted the model in three different ways: i) each of *n*_0_, *f_R_, c_R_*, and *d_T_* was allowed to be a patient-specific parameter (Model 1), ii) only *n*_0_ and *f_R_* were allowed to be patient-specific with *c_R_* and *d_T_* fixed at the mean values obtained in i) *(c_R_* = 0.78, *d_T_* = 0.14; Model 2), and iii) *c_R_* and *d_T_* were allowed to be patient-specific with *n*_0_ and *f_R_* fixed at the mean values obtained in i) (*n*_0_ = 0.59, *f_R_* = 0.04; Model 3). For 11 patients, Model 1 yielded only poor representations of the observed treatment dynamics (e.g. Patient 105 in Figure A6).

As the parameters associated with these fits are unlikely to be biologically meaningful we excluded these fits when computing the mean of the parameters taken forward to Models 2, and 3. For the same reasons, 11 patients were excluded in the analysis of Model 3 shown in Figure 6. Classification of patients into progressing and non-progressing was taken from the annotation provided in the data, where progression is defined as a series of three sequential increases of serum PSA > *4.0μg/L* despite castrate levels of serum testosterone. An overview of all fits for Models 1 and 3 is shown in Figure A6 and Figure A8, respectively. Fitting was done using the lmfit package in Python [32].

## 3. Results

### 3.1. Tumour control does not require resistance costs and is possible even if resistance arises in multiple locations

The aim of adaptive therapy is to delay disease progression by leveraging intra-tumoral competition. While it has long been thought that fitness costs associated with resistance are pivotal for the success of this strategy, our prior non-spatial work showed adaptive therapy can succeed even in the absence of a cost [22]. To test whether this prediction holds true if spatial interactions between cells are taken into account, we compare matched simulations in which the same tumour (identical parameters and random number seed) is treated once with continuous treatment and once adaptively. Progression is determined according to RECIST v1.1. as a 20% increase in tumour cell number relative to the start of treatment [33]. A representative example of our agent-based simulations is shown in Figures 2a & b. Despite the absence of a resistance cost and despite a random initial distribution of resistant cells, adaptive therapy is able to prolong TTP by about 15.3m (95% CI: [14.81m,15.86m]). Examination of the spatial dynamics in the simulation over time clearly illustrates competitive release of the pre-existing resistant cells (Figure 2b and Supplementary Movie 1). Of note, because of the random initial cell distribution the resistant population grows not as a single colony but emerges from multiple, independent “nests” simultaneously.

**Figure 2.**
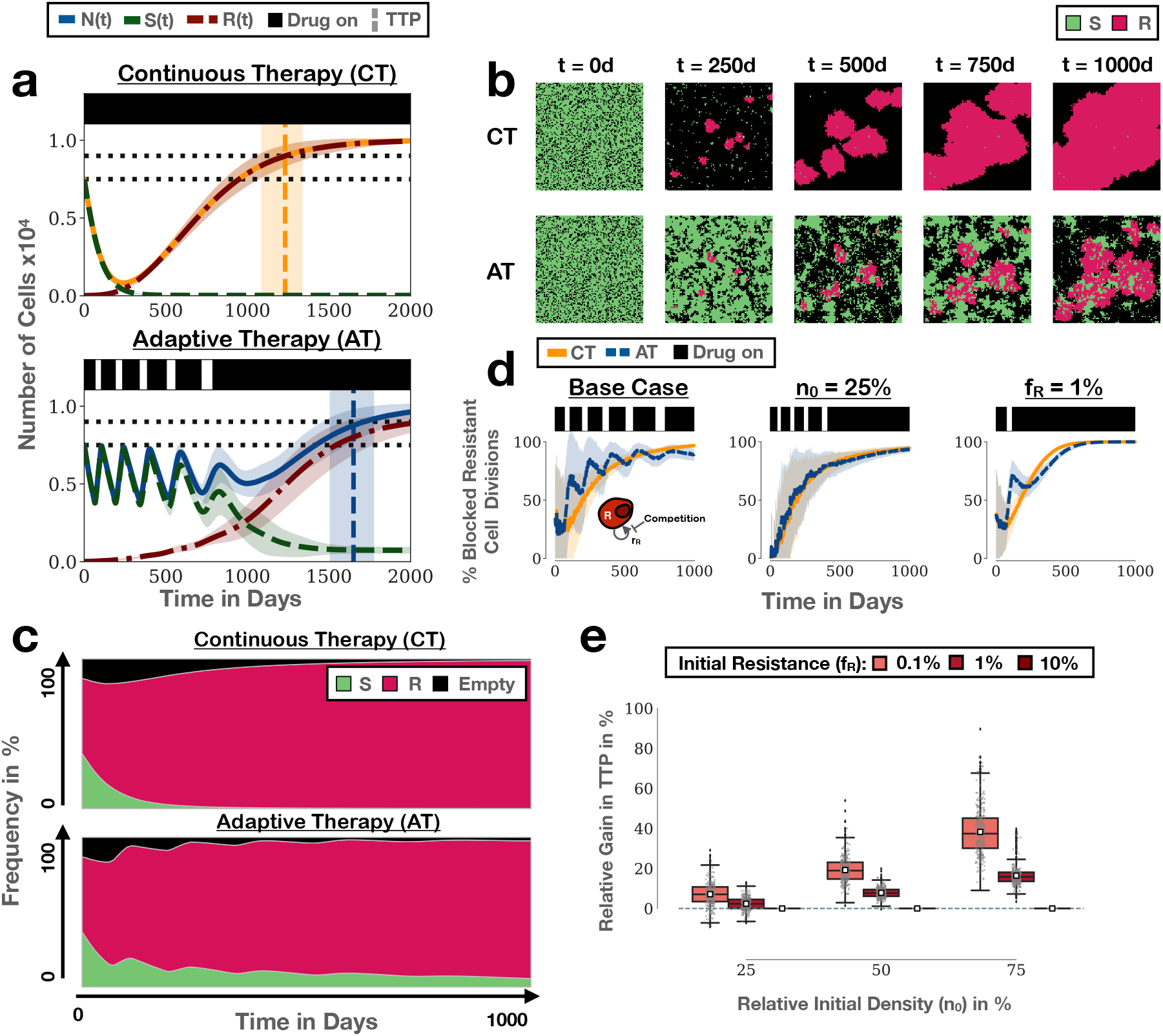
Adaptive therapy prolongs TTP by leveraging intra-tumoral competition and does not require a resistance cost. (**a**) Example of the treatment dynamics under continuous and adaptive therapy in our model ((*n*_0_, *f_R_*)=(75%,0.1%)). Shown are the mean and standard deviation (shading) of the tumour cell number (sensitive, resistant and total; 250 independent simulations). Black bars represents treatment. Horizontal dotted lines show the initial cell number and the cell number at progression, respectively. Vertical lines and associated shading mark the mean, and the 1^st^ and 3^rd^ quartile of the distribution of TTP. (**b**) Snapshots from one of the replicates in (a) illustrating how adaptive therapy delays competitive release of resistance. (**c**) Average frequency of sensitive cells, resistant cells, or empty space in a resistant cell’s neighbourhood in the simulations in (a). This shows there is not only inter- but also significant intra-specific competition during adaptive therapy. Values are the mean in the von Neumann neighbourhood across all resistant cells from 250 independent simulations. (**d**) Adaptive therapy (blue line) results in a higher proportion of blocked resistant cell divisions than continuous therapy (yellow line) in the initial stages of treatment in the simulations in (a) (left panel). However, the ability of adaptive therapy to induce blocking depends on the initial tumour cell density (centre panel; (*n*_0_, *f_R_*)=(25%,0.1%)) and the initial resistance fraction (right panel; (*n*_0_, *f_R_*)=(75%,1%)). Lines and shading indicate mean and standard deviation, respectively. (**e**) Gain in TTP achieved by adaptive therapy relative to continuous therapy as a function of *n*_0_ and *f_R_*. Adaptive therapy yields the greatest benefit when tumours are close to their carrying capacity and resistance is rare. Negative values denote cases in which continuous therapy achieves a longer TTP (1000 independent replicates per condition). Resistance costs and turnover are assumed to be 0 throughout this figure.

### 3.2. High tumour cell density and low initial resistance fraction maximise competition from sensitive cells

While competition is thought to be a driving mechanism behind adaptive therapy, it is challenging to quantify its role in real tumours. This is because it is difficult to rule out confounding factors, such as the effect of treatment de-escalation on tumour vasculature or the immune response [34,35], and because no simple biomarkers exist for measuring the strength of competition. Within our mathematical model we have precise control over the “experimental conditions” and we can measure quantities which are not accessible in real life. We seize this opportunity to examine how adaptive therapy modulates cell-cell competition for space by measuring the average composition of a resistant cell’s neighbourhood. This shows that the most frequent competitor of resistant cells are not sensitive cells but other resistant cells (Figure 2c). This is because as the resistant colonies become larger, most resistant cells are trapped in the centre of these colonies and are blocked by cells at the edge (Figure 2b). Accordingly, the proportion of blocked resistant cell divisions - a measure of the impact of competition - increases over time also under continuous therapy (Figure 2d; left panel). That being said, even though the proportion of resistant cells which directly compete with sensitive cells is small (Figure 2c), we see rapid and significant intensification of growth inhibition whenever treatment is withdrawn during adaptive therapy (Figure 2d; left panel). This is because when sensitive cells block growth at the edge of resistant colonies, this in turn also inhibits resistant cells in the centre (Figure 2b). As such, by leveraging both intra- and inter-specific competition adaptive therapy can maintain a higher level of growth inhibition during the initial phase of treatment, when resistance is still relatively rare, and can push back progression (Figure 2d; left panel).

Previous work by our group and others [15,16,19,22] has shown that the initial level of growth saturation of the tumour and the initial fraction of resistant cells are important determinants of the success of adaptive therapy. A parameter sweep over different values of *n*_0_ and *f_R_* confirms that the close proximity of the tumour to its carrying capacity and a small resistance fraction maximise the benefit of adaptive therapy also in the current model (Figure 2e). Moreover, consistent with this observation, we find that these conditions maximise the difference in the competition-mediated growth inhibition between continuous and adaptive therapy (Figure A3). We conclude that high growth saturation, due to close proximity to carrying capacity, and small initial resistance fractions are two factors that help to maximise the impact of inter-specific competition.

Moreover, we observe that *n*_0_ and *f_R_* have distinct effects on the number and impact of each adaptive therapy cycle, where we define a cycle by the time period between two subsequent crossings of the baseline tumour size, consisting of an on- and an off-treatment period. We find that when we reduce *n*_0_, the cycle is reduced only slightly, but the impact of each cycle on the reduction of the resistant cell growth rate is greatly diminished (Figure 2d; middle panel). Conversely, if we increase the initial resistance fraction, treatment fails after only a single cycle. However, this single cycle still induces blocking of resistant cells and results in a benefit in TTP (Figure 2d; right panel). This suggests that such tumours might be able to be controlled for longer using a less aggressive treatment regimen. Indeed, treatment of the same tumour with an adaptive protocol in which drug is withdrawn at a 30% reduction in size, increases the time gained from 2.5m to 6.5m (Figure A2).

### 3.3. In the absence of turnover, a resistance cost does not increase the relative benefit of adaptive therapy

One aim of this study was to investigate the role of resistance costs and cellular turnover on adaptive therapy. Figure 2a shows that neither costs nor turnover are necessary *per se* for adaptive therapy to be more beneficial than continuous therapy if the tumour is sufficiently close to its carrying capacity and resistance is rare. However, if these conditions are not met, then the benefit of adaptive therapy may be small, and in some cases longer control is, in fact, achieved with a continuous regimen (Figure 2e). This raises the question of whether in such cases a resistance cost may increase the benefit of adaptive therapy. We test this hypothesis for two different initial resistance levels in Figure 3a. The addition of a resistance cost slows down progression of the tumour under both protocols (Figure 3a), visible as a reduction in the size of resistant colonies (Figure 3b). An increased benefit of adaptive therapy is seen only for the case with the smaller value of *f_R_*, and this increase is modest (Figure 3a; right panel). Mapping out the impact of resistance costs in more detail corroborates this result (Figure 3c). We conclude that in the absence of turnover, resistance costs may increase the absolute time gained but will have little or no impact on the relative benefit of adaptive therapy. Only those patients in whom adaptive therapy would work also in the absence of a cost will benefit.

**Figure 3.**
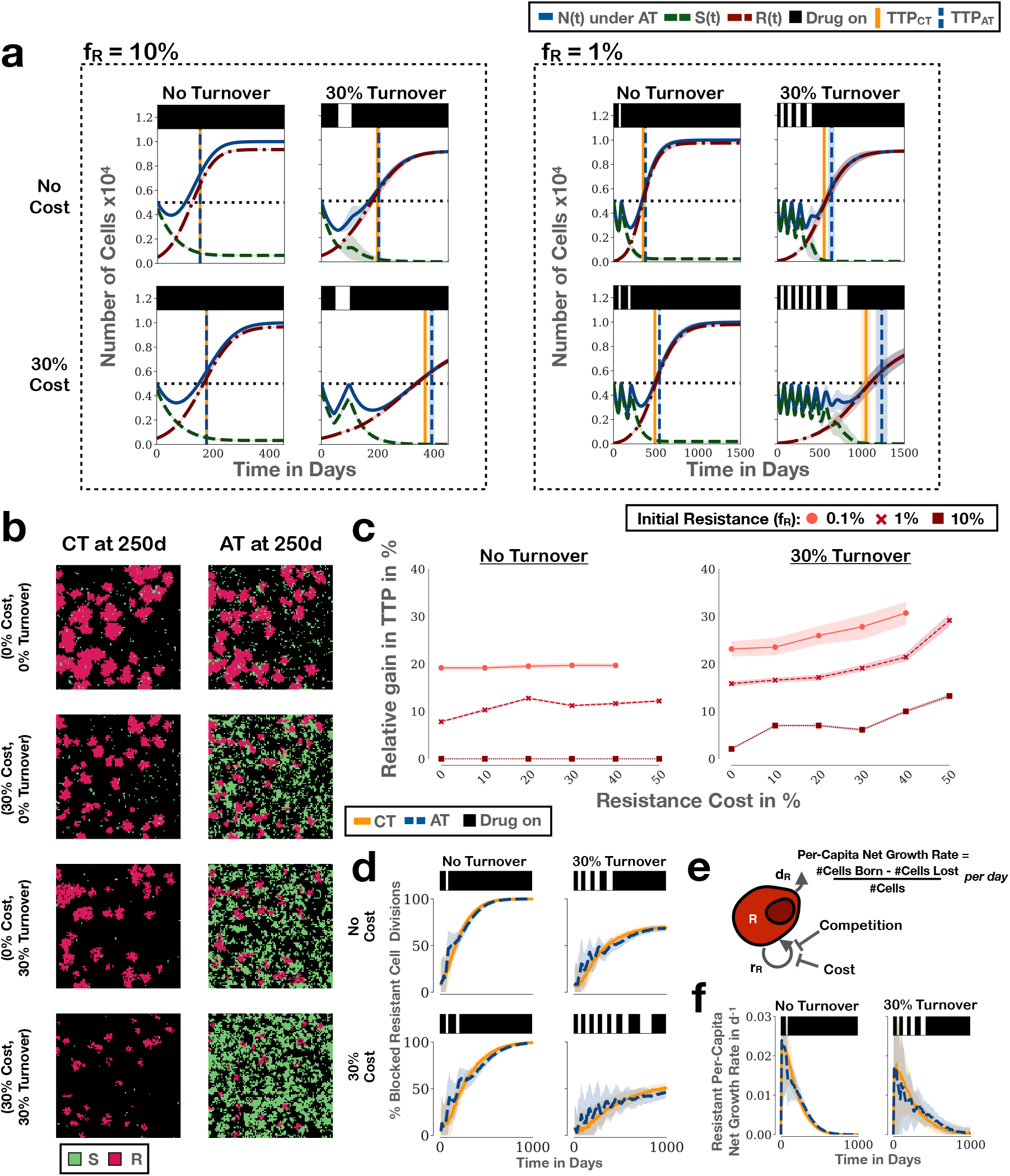
Resistance costs and high cellular turnover increase the benefit of adaptive therapy. Unless otherwise stated, lines and shading denote the mean and standard deviation of 250 independent replicates, respectively. (**a**) Simulations illustrating the role of resistance costs and turnover on the treatment response to adaptive therapy (*n*_0_ = 50%). Vertical lines and associated shading mark the mean, and the 1^st^ and 3^rd^ quartiles of the distribution of TTP. (**b**) Snapshots at time *t* = 250*d* from one of the replicates in the right hand panel in a) ((*n*_0_, *f*_R_) = (50%, 1%)). All 8 simulations were started from the same initial conditions and treated either continuously or adaptively assuming the indicated combination of cost and turnover. (**c**) Relative benefit of adaptive therapy over continuous therapy as a function of resistance cost, resistance fraction, and turnover (*n*_0_ = 50%). This illustrates that turnover modulates adaptive therapy and increases the impact of a resistance cost. Shown are the means (points) and the 95% confidence intervals (shading) for 1000 replicates per condition. (**d**) Impact of resistance costs and turnover on blocking of resistant cell division for the simulations in the right panel in (a). (**e**) TTP is determined by the per-capita net growth rate of resistant cells which is depends on both the birth and death rate of cells. Importantly, if death rates are high, then even moderate inhibition of cell proliferation by competition may result in large reductions in per-capita growth rate. (**f**) Per-capita growth rate of resistant cells as a function of time, illustrating how turnover helps to amplify the effects of competition on the resistant population’s net growth rate ((*n*_0_, *f_R_*) = (50%, 1%)).

### 3.4. Turnover increases the mean time gained by adaptive therapy by amplifying intra-tumoral competition

While cancer is known as a disease of increased cell proliferation, a large body of evidence suggests that cancers are also characterised by high rates of cell loss due to natural turnover, resource starvation and immune predation (e.g. [36-38]). We previously showed, in a non-spatial model, that high tumour cell turnover rates may increase the benefits of adaptive therapy [22]. To test whether this prediction holds true if spatial interactions are taken into account, we repeat our analysis from the previous section with a cell turnover rate of 30% relative to the proliferation rate, which is the value we previously estimated for prostate cancer [22]. For simplicity, we assume the same death rate for both sensitive and resistant cells and that dead cells are immediately removed from the domain. We find that the inclusion of cell death increases the average number of adaptive therapy cycles, and with it the benefit of adaptive therapy (Figure 3a). In addition, turnover amplifies the effect of resistance costs (Figure 3a & c).

To explain why turnover facilitates the control of the drug resistant population, we examine its impact on the tumour’s spatial architecture. This shows that addition of turnover results, not only in smaller, but also in fewer emerging resistant colonies, due to random extinction events (Figure 3b; see also Supplementary Movie 2). The presence of a resistance cost further amplifies this effect (Figure 3b). This shows that resistance costs and turnover not only reduce the growth rate of the tumour, but also change its spatial architecture.

Next we quantified the competition experienced by resistant cells in the presence, and absence, of turnover. We find that turnover reduces blocking of resistant cell divisions for both continuous and adaptive therapy (Figure 3d). This illustrates that tumour control depends, not only on the rate of proliferation of resistant cells, but also on the rate of cell death, as it is the per-capita *net* growth rate of resistance which determines TTP (Figure 3e). While increased turnover frees up space for cell division (see gaps at the centre of colonies in Figure 3b), it increases the impact of the blocking which does occur, as it limits the time, and so the opportunities, a resistant cell has for division. Indeed, comparing the per-capita growth rates of adaptive and continuous therapy we see that even though cell proliferation is less restricted with turnover, its impact on the per-capita proliferation rate is larger (Figure 3f). In summary, these results corroborate our hypothesis that the rate of tumour cell death is an important factor in adaptive therapy, and show how the impact of inhibition of cell proliferation depends on the rate of cell turnover.

### 3.5. The agent-based and corresponding ODE model yield quantitatively diverging predictions

Having found that the spatial model qualitatively recapitulates our prior non-spatial results [22], we asked whether the two models also agree quantitatively. To address this question, we compared simulations of the ODE and the agent-based version of our model assuming the same (exponential) rates for cell division, death and drug kill (Table 1; Figures 4a & b). This shows that while the dynamics agree qualitatively, there are important quantitative differences. Firstly, the ABM tends to predict longer TTP under both regimens (Figure 4a), and this discrepancy increases the higher the initial density and the smaller the initial resistance fraction (Figure 4b). At the same time, the cycling frequency is higher in the ABM with both short on- as well as off-times, resulting in a larger number of cycles (Figure 4a). Importantly, when we compare the relative benefit of adaptive therapy predicted by the two models we find that the ABM tends to forecast a smaller gain than the ODE model, especially if turnover is included in the model (Figures 4b & c). In addition, there is significant variation in the possible outcomes in the ABM, with some patients even progressing faster on adaptive than on continuous therapy - an outcome not possible in the ODE model (Figure 4c; for a proof of the latter see [16,39]). This indicates that, while the simple ODE model is useful for building qualitative understanding of adaptive therapy, spatial and stochastic effects may significantly alter the quantitative dynamics actually observed in the wet-lab or the clinic.

**Figure 4.**
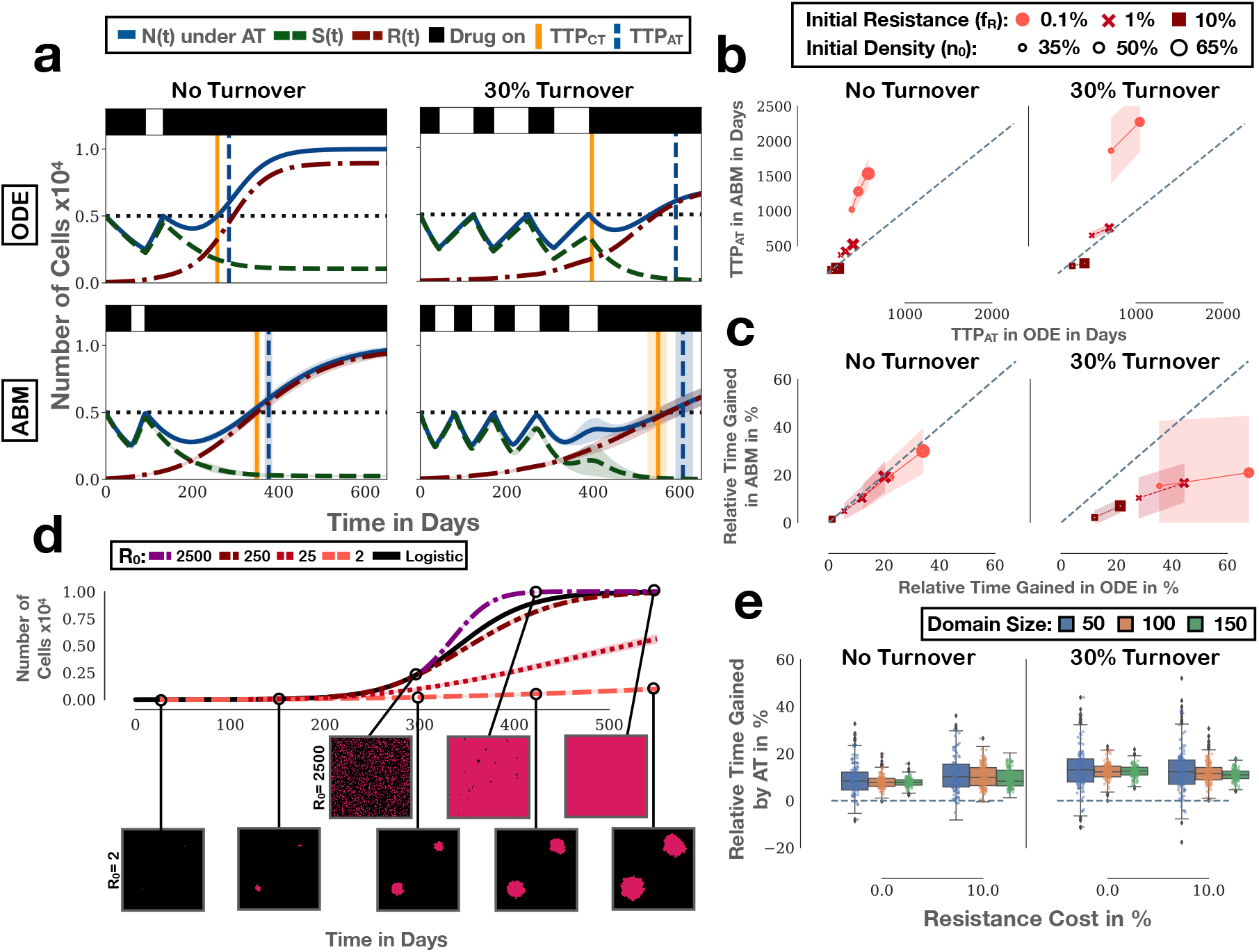
Comparing the spatial (ABM) with the non-spatial (ODE) model from [22]. Lines and shading denote the mean and standard deviation, respectively. (**a**) ODE and ABM simulations with the same parameters ((*n*_0_, *f_R_*, *c_R_*) = (50%,1%,0%); 250 replicates for the ABM). (**b**) Comparison of the TTP under adaptive therapy in the ODE and ABM (c_R_ = 0%). The ABM predicts faster progression when initial resistant cell numbers are high, and slower progression when numbers are low (*n* = 1000 replicates of the ABM). In the presence of turnover, tumours starting from *n*_0_ = 65% can not grow above the threshold defining progression and thus, no TTP can be obtained. (**c**) Comparison of the relative benefit of adaptive therapy ((TTPA_AT_-TTP_CT_)/TTP_CT_) in the two models. The ODE model tends to predict larger benefit than the ABM. Again, no TTP can be obtained for *n*_0_ = 65%. (**d**) Depending on the initial cell number (*R*_0_) the resistant population will grow sub- or super-logistically (logistic growth: black solid line). Resistant cells were seeded and grown in isolation and without drug (*n* = 250). To allow direct comparison with logistic growth, the ABM curves were shifted to start at the time at which the logistic model reached the corresponding starting number of cells. (**e**) Domain size (l) of the ABM reduces the variance in outcomes further corroborating the importance of the initial resistant cell number ((*n*_0_, *f_R_*) = (50%, 1%); *n* = 1000 independent replicates).

### 3.6. The number of independent resistant nests is a driver of progression dynamics

To understand why the discrepancy between ABM and ODE model arises, we examined the growth dynamics in both models in more detail. We find that when the initial resistance fraction is small, the resistant population in the ABM expands more slowly than in the ODE model, but the converse holds true when the initial resistance fraction is large (Figure A4). To explain this, we simulated the resistant population in isolation, starting from different initial cell numbers. Our results demonstrate that different initial cell numbers generate distinct growth kinetics (Figure 4d). When initiated from two cells, the resistant population expands as two colonies and grows much more slowly than predicted by a logistic ODE model, as most cells are trapped by their neighbours (Figure 4d). As the number of cells, and so the number of independent nests and the surface to volume ratio, is increased, the growth of the population speeds up until it exceeds that of logistic growth (Figure 4d). This explains the observed difference in the predicted TTP in Figure 4b, and why this discrepancy worsens for smaller resistance fractions.

### 3.7. The impact of turnover is reduced by the neighbourhood size

Similarly, we can explain the reduced impact of turnover on TTP by differences in cell growth dynamics between the two models. Turnover has two effects: On one side, it limits a cell’s lifespan and so the number of opportunities it has to divide. The higher turnover, the smaller is this number, and the greater the impact if a division is blocked by a competitor [22]. On the other hand, death of a neighbour opens up space for cell division which partially off-sets the cell loss caused by turnover (Figure 3d). Importantly, in the ABM each cell can divide into four potential sites which allows for more divisions to take place at high cell density than predicted by the ODE model, which is why the impact of turnover is reduced (Figure A5). Repeating this comparison with a Moore neighbourhood, in which a cell can divide into all 8 surrounding sites, diminishes the impact of turnover even further (not shown). We conclude that spatial constraints on tumour growth are an important factor in determining the impact of cell turnover.

Turnover also causes larger variability in treatment outcomes in the ABM (Figure 4b). This is because turnover can make individual nests go extinct, which changes the growth dynamics of the tumour, and can even result in the complete eradication of the resistant population (see also Section 3.8). That being said, this is a stochastic effect which is reduced when we increase the total number of cells in the simulation (Figure 4e). Furthermore we find that also the frequency of cases in which adaptive therapy performs worse than continuous therapy decreases with increasing domain size (Figure 4e). This indicates that failure of adaptive therapy in the ABM is mostly driven by stochastic events. We conclude that a further effect of turnover is to amplify the impact of stochastic events on treatment outcome, especially when initial resistant cell numbers are small.

### 3.8. The number of locations in which resistance arises is an important factor in adaptive therapy

An important implication of these results is that the spatial distribution of resistant cells within the tumour impacts treatment response. The existence of multiple resistance mechanisms, and the increasing evidence for the role of phenotypic plasticity and environmental factors, suggest that resistance may develop in multiple locations in parallel [40]. Evidence for this has been observed, for example, in colorectal cancer [41], lung cancer [42], and melanoma [43]. To investigate how adaptive therapy is affected by whether resistance arises at a single site or in multiple locations we conducted a series of experiments in which we seeded the same number of resistant cells, either as a single cluster, or as a set of two clusters, at varying distances apart (Figure 5a; top row). We placed eight resistant cells as a 2 × 4 rectangle in the centre of the domain. Subsequently, we compared this scenario to those in which we split this cluster into two nests of size 2 × 2 which we placed at varying distances apart from each other. Sensitive cells were seeded randomly in the domain to achieve a total initial density of *n*_0_ *=* 50%.

**Figure 5.**
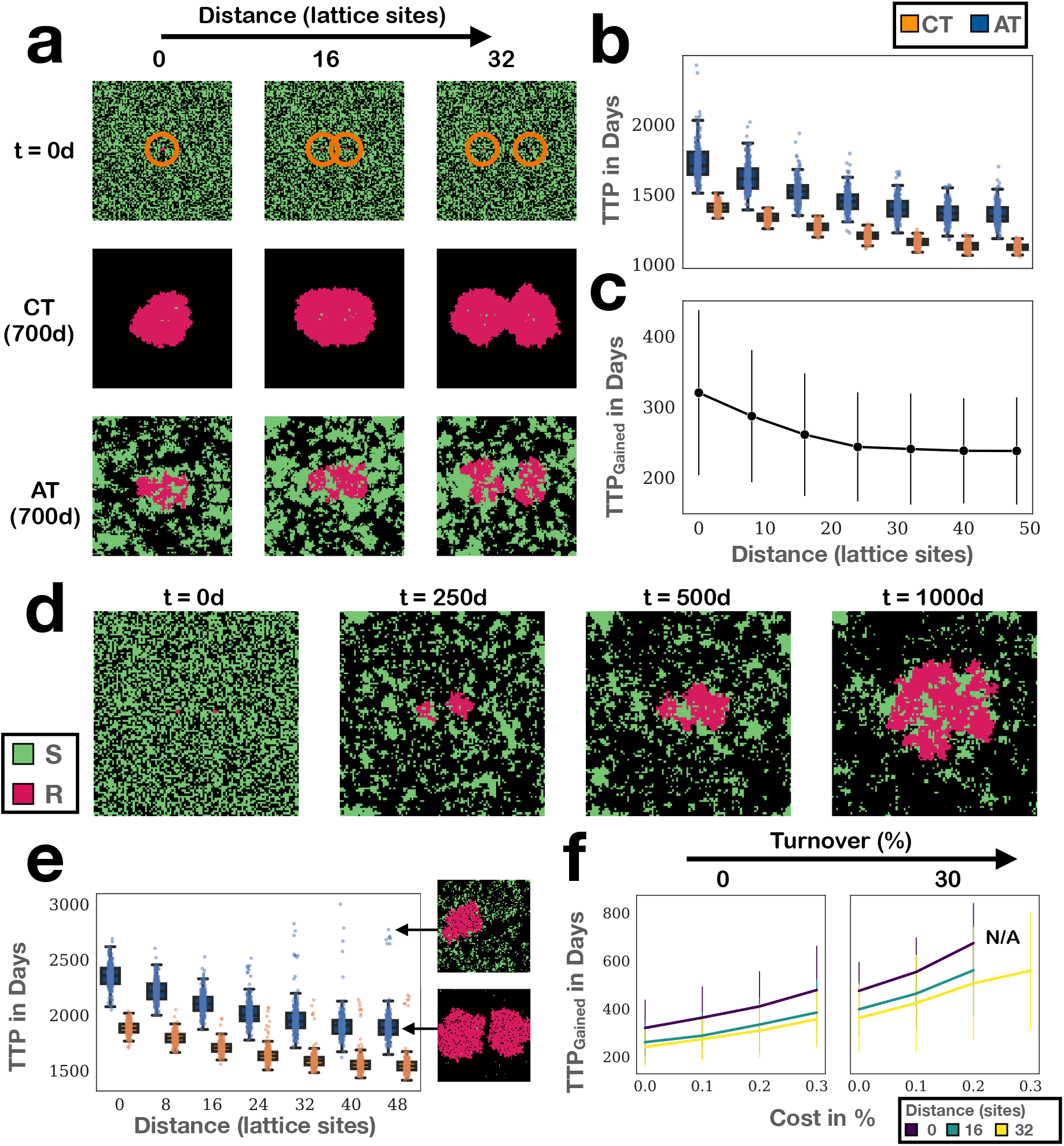
As a result of parallel mutation events, or because of mediation via environmental factors, resistance can arise in multiple locations at once. This poses a number of challenges for resistance management. (**a**) To investigate the impact of the initial spatial distribution of resistance on treatment response we seeded eight resistant cells either as a cluster in the centre, or as two nests at varying distances apart (marked with orange rings in the top row). Shown are representative snapshots from simulations for three different initial levels of separation ((*n*_0_, *c_R_*, *d_T_*) = (50%,0%,0%)). (**b**) TTP of continuous and adaptive therapy decreases as the separation between the nests increases (1000 replicates per conditions; parameters as in (a)). (**c**) Similarly, the benefit of adaptive therapy decreases the more separated are the nests (1000 replicates per conditions; parameters as in (a)). Error bars mark one standard deviation. (**d**) Representative snapshot from a simulation in (a) in which the nests are initially 16 lattice sites apart, which illustrate why control of multiple nests is more challenging. While the left nest can initially be controlled, the right nest escapes and, in turn, triggers release of the left nest. (**e**) TTP as function of separation in the presence of turnover (*d_T_* = 30%). Turnover can cause extinction of one of the two nests which significantly increases TTP (1000 replicates per condition). Insets show illustrative snapshots at *t* = 1500*d*. (**f**) Effect of the initial spatial distribution of resistance on the relationship between cost, turnover and gain of adaptive therapy (1000 replicates per condition). Error bars show one standard deviation.

Our results show that resistance arising in multiple locations causes faster progression and reduces the benefit of adaptive therapy. The further the nests are seeded apart from each other, the quicker resistant cells expand because they are competing with sensitive cells which are killed by drug, and not with other resistant cells (Figure 5a). This results in shorter TTP (Figure 5b) and a smaller benefit of adaptive therapy (Figure 5c). In addition, expansion of one resistant nest can also support the growth of another: While the left of the two nests in Figure 5d is initially constrained by sensitive cells (*t* = 250*d*), the right nest is not, and is able to expand (*t* = 500*d*). This triggers more and more treatment which eventually results in the competitive release also of the left nest (*t* = 1000*d*). This indicates that competition of resistant cells with each other is an important, but overlooked, factor in adaptive therapy. While there is more competition with sensitive cells when the resistant nests are seeded apart, better control is achieved when they are clustered together because it allows adaptive therapy to leverage both inter- as well as intra-specific competition.

### 3.9. The spatial distribution of resistance modulates the impact of cost and turnover

Next, we studied how space modifies the impact of resistance costs and turnover. We find that also in the presence of turnover mean TTP and the benefit of adaptive therapy decrease with increasing nest separation (Figure 5e). At the same time, turnover can cause the random extinction of one of the two nests, or both, which significantly extends TTP and results in a high variability in outcomes, especially if the separation between nests is large (insets in Figure 5e). In addition, greater spread of resistant cells reduces the benefit adaptive therapy can derive from resistance cost and turnover (Figure 5f). We conclude that the spatial distribution of resistance plays a significant role in the response to adaptive therapy, and its potential benefit.

### 3.10. The cycling frequency of patients undergoing intermittent androgen deprivation therapy may reflect different distributions of resistance

Androgen deprivation therapy is an integral part of prostate cancer treatment as many tumours are initially dependent on androgen signalling for their growth [44]. The inevitable development of androgen-independence, as well as the impact of treatment on quality of life, has motivated a number of intermittent therapy trials in prostate cancer (e.g. [23,45,46]). The trialled algorithms administer treatment until the levels of prostate-specific antigen (PSA), a blood-based biomarker used for tracking tumour burden, are reduced to normal levels. Subsequently, treatment is withdrawn until PSA levels again exceed some upper limit, when treatment is re-instated. Interestingly, it is observed that some patients are cycling rapidly under treatment whereas others are cycling more slowly (Figure 6a), although the relationship of cycling frequency and outcome is unclear [23].

**Figure 6.**
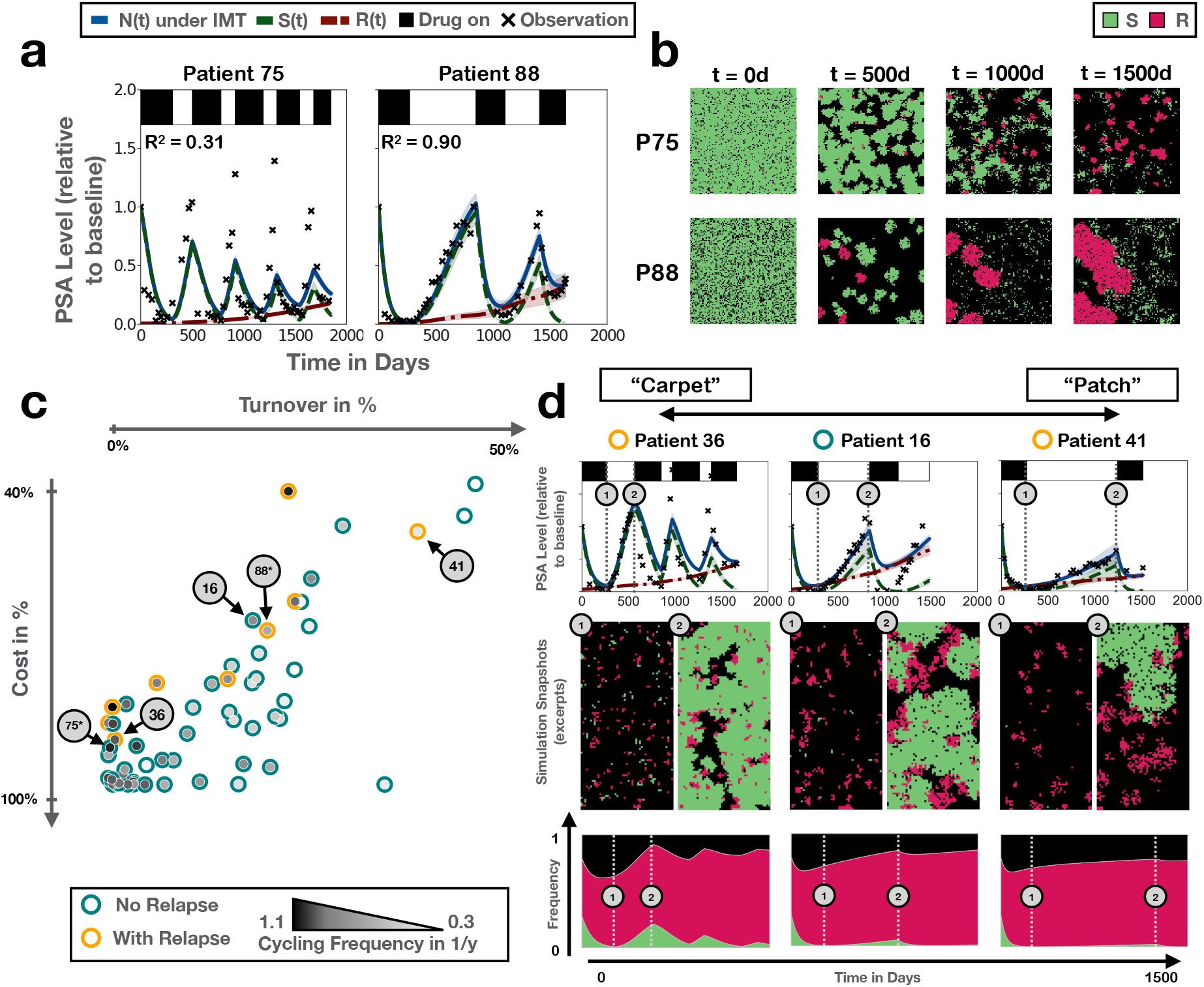
Analysis of the cycling dynamics of 67 prostate cancer patients undergoing intermittent androgen deprivation therapy in the trial by Bruchovsky et al [23]. Lines and shading denote the mean and standard deviation of 250 independent replicates, respectively. (**a**) Representative fits of the ABM to the PSA data from a fast and a slow cycling patient (Patients 75 and 88, respectively). (**b**) Snapshots from one of the simulations in (a), showing the distinct patterns of resistance growth in the two patients. (**c**) Negative correlation between the estimated values of cost and turnover, revealed when fitting just cost and turnover on a patient-specific basis. Note that while Patients 75 and 88 are marked for reference, their fits here are not those shown in (a) which were fitted with all four parameters being allowed to vary. For an overview of all fits for this 2-parameter model, see Figure A8. (**d**) Treatment trajectories, simulation snapshots and neighbourhood composition, illustrating the dynamics for patients in different areas of the parameter space in (c) ranging from what we term “carpet”-like appearance (many small, resistant colonies) to “patch”-like appearance (few, but large, resistant colonies; see also Supplementary Movie 4).

Using the ODE model (Equations (2)-(4)) we previously showed that we can explain the different cycling dynamics across patients in the Phase II study by Bruchovsky et al [23] by a combination of the tumour turnover and the resistance cost, suggestive of different underlying disease biology [22]. In order to test for possible differences in the tumours’ spatial architectures, we fitted the ABM to these same data, which consist of monthly PSA measurements of 67 patients undergoing intermittent androgen deprivation treatment for recurrent, locally advanced prostate cancer. We allowed the values of *n*_0_, *f_R_*, *cR*, and *dT* to vary on a patient-specific basis, whilst keeping all other parameters fixed across the cohort at the values shown in Table 1. Given the stochastic nature of our simulations we assembled the fit for each patient from the mean of 25 independent stochastic replicates. We find that, despite the simplicity of the model, it can recapitulate the cycling dynamics for a majority of patients, and can describe fast, as well as slowly, cycling patients (Figure 6a; see also Figure A6).

Moreover, fast and slow cyclers are associated with distinct spatial dynamics in our simulations (Figure 6b and Supplementary Movie 3). In fast cycling patients, the tumour has a “carpet-like” structure with many small, independent, sensitive and resistant nests (Patient 75). In contrast, in slowly cycling patients growth is driven by only a handful of large, “patch-like”, colonies (Patient 88). In order to understand which parameters are key in driving this behaviour, we fitted the model keeping either the initial conditions (n_0_ and f_R_) or the cell kinetic parameters (c_R_ and *d_T_*) fixed across the cohort. We find that allowing just cost and turnover to be patient-specific can explain the data almost as well as the full four parameter model (Figure A7). In contrast, the model assuming that inter-patient variability is caused by different initial conditions fits poorly (Figures A7 & A8). In addition, these results reveal a negative correlation between the fitted values for the resistance cost and turnover (Pearson’s correlation coefficient: *r* = −0.76, p < 0.01), with fast cyclers being associated with large values of cost and small values of turnover, and vice versa for slow cyclers (Figure 6c). Moreover, the differences in the spatial architecture persist and we can now explain why (Figure 6d and Supplementary Movie 4): When turnover is low and cost is high (fast cyclers), then most resistant cells present at the start of treatment will survive, but only expand very slowly, which yields the diffuse, carpet-like appearance of these tumours. In contrast, when turnover is high and cost low (slow cyclers), then many initially present resistant colonies will go extinct, but those that do survive will be able to expand more rapidly. Thus, the patch-like appearance of these tumours.

A further interesting observation is that no patients are to be found on the top left corner of the graph, and only one patient is located on the bottom right corner (Figure 6c). This can be explained by the trial’s patient selection criteria. When cost and turnover are small, response is weak so that patients in this parameter regime would not have been able to produce the initial PSA normalisation required for study inclusion. Conversely, patients in whom both cost and turnover are high will show durable responses and are, thus, unlikely to be refractory after initial therapy (recall Figure 3a for how cost and turnover impact response). Moreover, previously, we observed that patients who progressed on the trial were characterised not by a lack of cost or turnover but by a smaller combination of the two [22]. Our current analysis corroborates this context-dependence of the resistance costs. Progressors (yellow crosses) cluster along the upper boundary of the line of fits (Figure 6c). Accordingly, we detect no statistically significant difference in the turnover estimate (Mann-Whitney test, *U* = 210, *p* = 0.5), but a significant difference in the sum of the two *(c_R_* + *d_T_*; Mann-Whitney test, *U* = 45, *p* < 0.01). That being said, we do observe a statistically significant difference in the estimated cost values (Mann-Whitney test, *U* = 95, *p* < 0.01).

To further characterise the differences between the inferred spatial architectures of fast and slow cycling patients, we quantified intra-tumoral competition. This shows that in slowly cycling patients almost all competition is intra-specific, whereas in fast cyclers competition with sensitive cells plays more of a role (Figure 6d). Overall, this supports our hypothesis that different cycling speeds reflect different underlying disease biology [22], and suggests this may not only be driven by differences in the cell kinetics but also manifest itself in distinct tumour architectures and competition landscapes.

## 4. Discussion

Evolutionary theory has developed into a unifying framework through which to understand cancer initiation, progression, and treatment response [47]. This has triggered the development of treatment strategies which aim to steer tumour evolution rather than ignore it [6,48-50]. Adaptive therapy is such an approach and seeks to control drug resistance by exploiting intra-tumoral competition between treatment sensitive and resistant cells, in a patient-specific manner.

The aim of this study was to better understand the impact of a tumour’s spatial organisation on adaptive therapy. To do so, we studied a simple 2-D, on-lattice ABM in which tumour cells are classified as either drug-sensitive or resistant. This shows that adaptive therapy is, on average, superior to continuous therapy under a wide range of parameters, assuming a cure is not possible and recurrence is primarily driven by pre-existing resistance. Furthermore high initial tumour cell density, and low initial resistance fractions, maximise the benefit of adaptive therapy, and even in the absence of a resistance cost we are able to slow the emergence of resistance by leveraging competition, indicating that such a cost is not a requirement for adaptive therapy. Overall, these results are consistent with the conclusions we and others have drawn in previous non-spatial [11,15,16,18,22,51] and spatial theoretical studies [11,19], and with experimental evidence in cancer [11] and bacteria [52]. This is an important finding as it indicates that conclusions drawn from simple, non-spatial models appear to carry over qualitatively to spatially growing tumours.

That being said, when we perform a more detailed comparison of the ABM with its corresponding non-spatial ODE model, we observe significant quantitative differences between the models. In particular, while adaptive therapy is superior in both models, its relative benefit compared to continuous therapy is smaller in the ABM, and there is less gain from the presence of resistance costs and turnover. This is because the two models make different assumptions about spatial competition. In the ODE model, cells are assumed to be perfectly mixed, so that all cells experience the same competitive growth inhibition, which is simply a linear function of the total cell density. In contrast, the lack of migration in the ABM results in spatial segregation of different colonies, so that the competitive inhibition experienced by a cell depends on the cell’s local neighbourhood, and varies across the tumour. Consequently, sensitive cells in the ODE model will always be able to competitively suppress resistant cells, whereas in the ABM this is only possible if the sensitive colony grows in close vicinity to the resistant colony. This indicates that the tumour’s spatial organisation is likely to be an important factor in adaptive therapy. Moreover, it shows that a detailed understanding of intra-tumoral competition is required, in order to determine whether or not a patient will receive a clinically meaningful gain from adaptive therapy. This point is also supported by the recent work by Viossat and Noble [16], and Farrokhian et al [53], who found that while different ODE models of adaptive therapy agree qualitatively, there are significant differences in their quantitative predictions depending on how competition is modelled.

In order to gain an understanding of intra-tumoral competition during treatment, we explicitly quantified it in our simulations. To the best of our knowledge, this is the first time the effect of adaptive therapy on intra-tumoral competition has been explicitly quantified. Our results illustrate how treatment breaks leverage inter-specific competition with sensitive cells to slow the expansion of pre-existing resistant populations. Furthermore, we show that intra-specific competition of resistant cells with each other is an important, but so far overlooked, factor in adaptive therapy. As a resistant cell expands, most of its daughters will end up trapped in the central core of the resistant colony, able to divide only upon the death of another resistant cell (or upon migration out of the centre). As such, adaptive therapy is most effective when resistance is clustered in a single location, and surrounded by sensitive cells, because it can leverage inter-specific competition at the edge of the resistant colony to maximise intra-specific competition between resistant cells at the core (Figure 7a).

**Figure 7.**
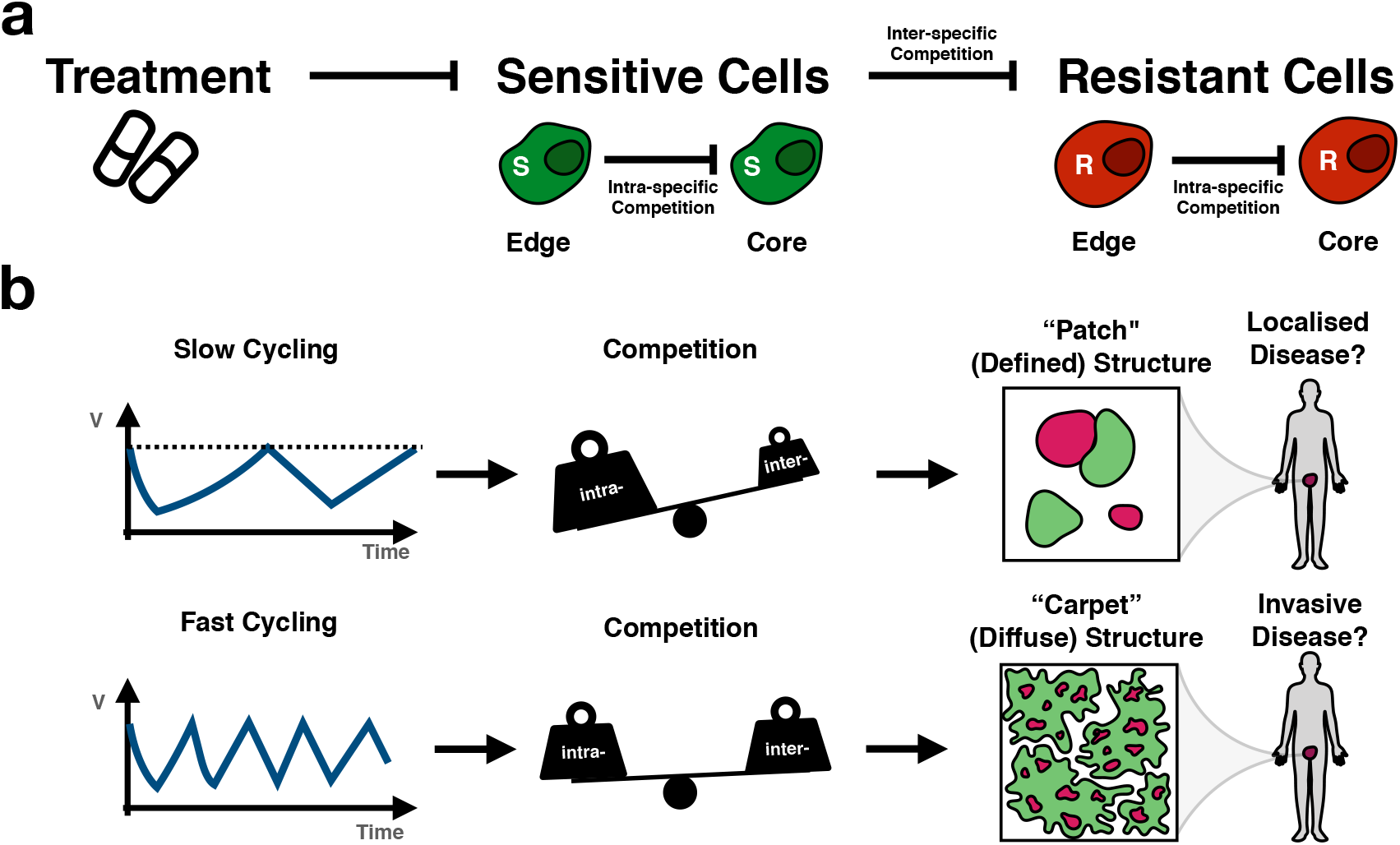
The tumour’s spatial architecture determines the ratio of intra- to inter-specific competition, and thereby the response dynamics under adaptive therapy. (**a**) The current paradigm foresees that adaptive therapy keeps resistant cells in check by inter-specific competition from sensitive cells, which can be controlled via treatment ([6,8]; adapted from [22]). Here we show that intra-specific competition within each population is a further important factor. (**b**) Our analysis suggests that the more diffusely growing a tumour, the greater the fraction of inter-specific competition. In addition, this may be reflected in faster cycling frequency under intermittent treatment. Going forward, we propose that incorporating knowledge of the tumour’s spatial architecture and resulting competition structure will help to design more effective adaptive therapy strategies.

An important implication of this observation is that it matters how resistance is distributed across the tumour. If resistance arises in a single location it can be controlled more effectively with adaptive therapy than if resistance is present at multiple sites, either within the same lesion, or at different metastatic sites within the body. As such, we generalise previous results by Bacevic et al [11] who showed in a computational model that resistant cells arising near the surface of a spheroid are harder to control than resistant cells arising near its core, and we characterise in detail the non-trivial relationship between a tumour’s spatial organisation and adaptive therapy.

But how may we infer the spatial distribution of resistance? Tissue biopsies would provide the most direct and detailed measurements, but are invasive and often impractical. As an alternative, we show that it may be possible to use mathematical modelling to gather spatial insights from the patient’s longitudinal response dynamics (Figure 7b). We fit our ABM to PSA data from prostate cancer patients undergoing intermittent androgen deprivation therapy. We find that the speed at which patients cycle between treatment on- and off-periods correlates with distinct spatial organisations of the tumours in our simulations. While experimental validation of this hypothesis is outstanding, we do note that Bruchovsky et al [23] reported a “suggestive trend that a Gleason score <6 may be associated with a slightly longer time off treatment in the initial 2 cycles”. Indeed, lower Gleason scores indicate a more defined tissue architecture, which would be consistent with the more patch-like structure our model predicts for slowly cycling patients. As such, this hypothesis warrants further investigation.

In addition, it may also be possible to gather some information about the spatial distribution of resistance from the characteristics of the resistant population. In particular, if resistance is driven by a single clone, then it will likely be initially confined to a single, or at most a small number of, sites within the tumour. In contrast, if resistance is driven by multiple clones, as has, for example been observed in colorectal cancer [41], then it is likely to exist in multiple locations simultaneously. Liquid biopsies are showing promise at detecting and characterising the clonality of emerging drug resistance, and as such may provide a useful tool for informing adaptive therapies [40,41].

While we cannot easily alter it, understanding the spatial distribution of resistance in a patient may be relevant in the design of adaptive treatment schedules. Gallaher et al [19] found in an off-lattice ABM that the rate of cell migration (and therefore of spatial mixing) determined whether a modulation-based adaptive algorithm (treatment is modulated in small increments, rather than withdrawn), or a vacation-based algorithm (treatment is either on or off) was more effective. In particular, spatially confined tumours favoured modulation, whereas in invasive tumours more benefit was derived from the vacation-based strategy [19]. Consistent with this conclusion, Yu et al [54] found in a Phase I trial in EGFR-mutant lung cancer that a carefully designed combination of low and high doses of erlotinib controlled central nervous system metastases more effectively than standard-of-care continuous dosing. Conversely, Benzekry and Hahnfeldt [55] concluded from the study of an ODE model that metronomic treatment (low dose, high frequency) may be more effective in controlling metastatic disease than aggressive standard-of-care treatment (high dose, low frequency). Investigating how to best adapt treatment when there are multiple resistant nests, and/or metastasis, is an important direction of future research.

A final observation we make is that there can be significant variation in the benefit of adaptive therapy between stochastic replicates despite identical model parameters. Variance depends on the number of resistant cells initially present in the simulations, and their distribution, which further highlights the importance of the spatial distribution of drug resistance within the tumour. Moreover, even though resistance costs and turnover increase the average benefit of adaptive therapy they also increase variability in outcomes. As a result, the greater the average benefit of adaptive therapy the more likely we are to see a patient fair much better or worse than expected. While the cell numbers in our simulations are unrealistically small, and as such our simulations are not suited to make quantitative statements about the magnitude of this problem, Hansen et al [56] have recently raised similar concerns. We, thus, advocate further study of the impact of inter-patient variability in adaptive therapy, using, for example, Phase i trials [57], in order to inform future clinical trial design.

In aiming to keep our model tractable we have made a number of simplifying assumptions. We assume no movement and no pushing of cells, which has been shown by Gallaher et al [19] to reduce the benefit of AT, as it allows resistant cells to squeeze through surrounding sensitive cells. Moreover, for computational reasons, we restricted our analysis to a 2-D setting, which is arguably more representative of *in vitro* cell culture than a 3-D human tumour. We hypothesise that the extra dimension will hinder tumour control as it will allow resistant cells to more easily find space into which they can divide. That being said, we have also neglected the role of non-tumour tissue, which acts as an additional competitor for space and resources in the tumour, and may help to control resistant subpopulations [18]. We, and others, have also investigated the important role of metabolism in regulating tumour progression, immune dynamics and treatment response [58-60]. The role of the tumour intrinsic metabolism versus the extrinsic tissue microenvironment metabolism has also not been considered here. However, their inclusion is more likely to enhance the impact of adaptive therapy than diminish it due to the potential for increasing the cost of resistance (e.g. [22,61,62]).

A final, important caveat is that we have not explicitly modelled the mechanism by which resistance arises. Depending on whether it arises through mutation, phenotypic switching, or is environmentally induced, this may drive different initial distributions of resistant cells, and will also result in different dynamics during treatment due to *de novo* resistance acquisition (see also [16,19,51,63]). Furthermore, our model cannot explain how some of the initial tumour compositions we have analysed would have arisen prior to treatment. For example, if resistance costs and turnover are assumed to be high, then resistance will disappear if the tumour is left untreated (not shown).

To sum up, in this paper, we have consolidated and advanced our understanding of how competition between tumour cells may be leveraged by careful treatment modulation. We have shown that the spatial organisation of resistant populations is an important, and understudied factor in cancer treatment. This strengthens the argument for patient-specific, adaptive therapy protocols that explicitly consider not only a tumour’s evolution but also its ecology.

## Author Contributions

Conceptualization, M.S., P.M. and A.A.; methodology, M.S.; software, M.S.; formal analysis, M.S.; investigation, M.S.; resources, P.M. and A.A.; data curation, M.S.; writing-original draft preparation, M.S.; writing-review and editing, P.M. and A.A.; visualization, M.S.; supervision, P.M. and A.A.; project administration, P.M. and A.A.; funding acquisition, P.M. and A.A. All authors have read and agreed to the published version of the manuscript.

## Acknowledgments

We thank Ruth Baker, and Gregory Kimmel for helpful discussions about the implementation of the stochastic simulation algorithm.

## Abbreviations

The following abbreviations are used in this manuscript:

ABM: Agent-based model
ODE: Ordinary differential equation
TTP: Time to progression

## Appendix A. Stochastic simulation algorithm

A flow-chart of our model is shown in Figure 1b. We implement this model using the following fixed time-step stochastic simulation algorithm:

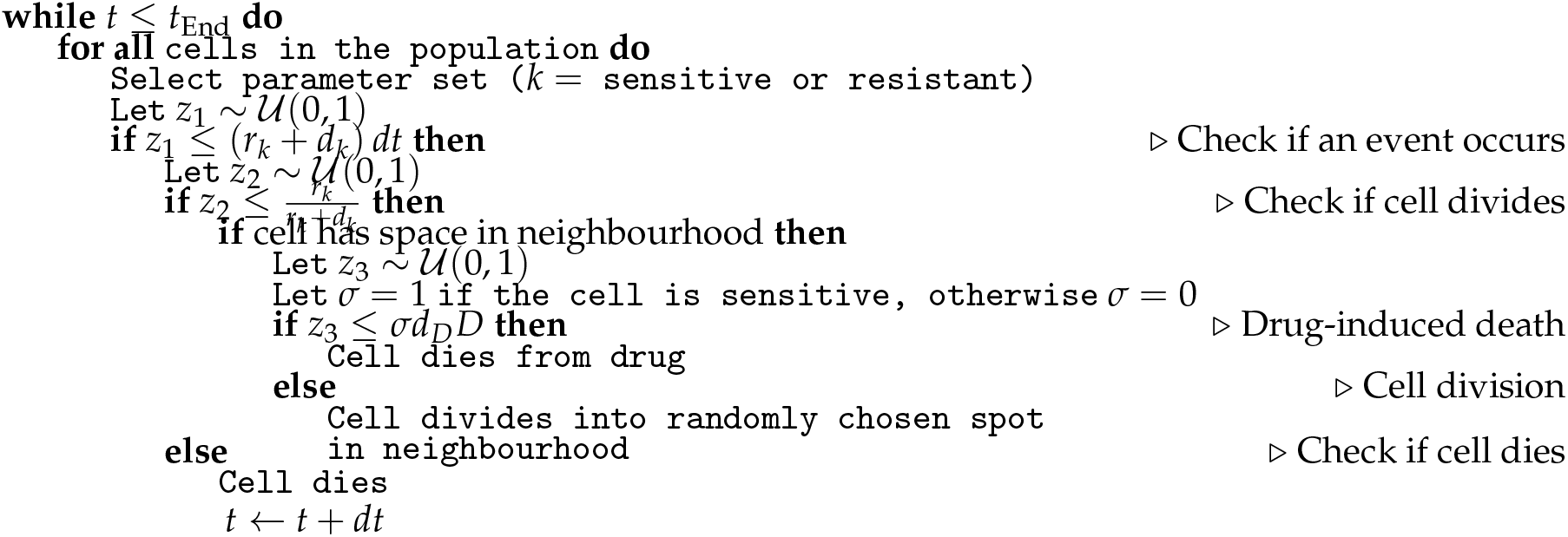

Here, 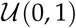 denotes the uniform distribution on [0,1], and t_End_ the end time of the simulation in days.

## Appendix B. Consistency analysis

A consistency analysis serves to determine the number of independent replicates required to obtain a sample of outcomes representative of the stochastic process. We adopt the protocol from [25] and apply it to corner cases of our parameter space, (*n*_0_, *f_R_*, *c_R_*, *d_T_*). The idea is to choose a sample size, *n*, obtain *k* independent samples of this size, and compare the *k* distribution of model outcomes. The aim is to find a value of *n* so that the *k* distributions are sufficiently similar and any one of them is representative of the others. We computed the difference in TTP for adaptive and continuous therapy for 10 samples of sample sizes *n* = 10,50,100,250,500,1000, and 1500, respectively (68,200 simulations total). In Figure A1a we illustrate, for one parameter combination, how the thus obtained 10 outcome distributions for each value of *n* become almost indistinguishable for *n* ≥ 250. To quantify consistency we measured the mean value of each distribution and the proportion of runs in which adaptive therapy performed worse than continuous therapy. This corroborates that *n* ≥ 250 generates outcome distributions with very similar mean values and lower tails for a range of parameter combinations (Figure A1b & c). However, some parameter sets converge more slowly than others. Thus, we choose a value of *n* = 1000 for all analyses except for time-series plots (e.g. Figure 2a & b) for which we choose *n* = 250 for computational reasons.

**Figure A1.**
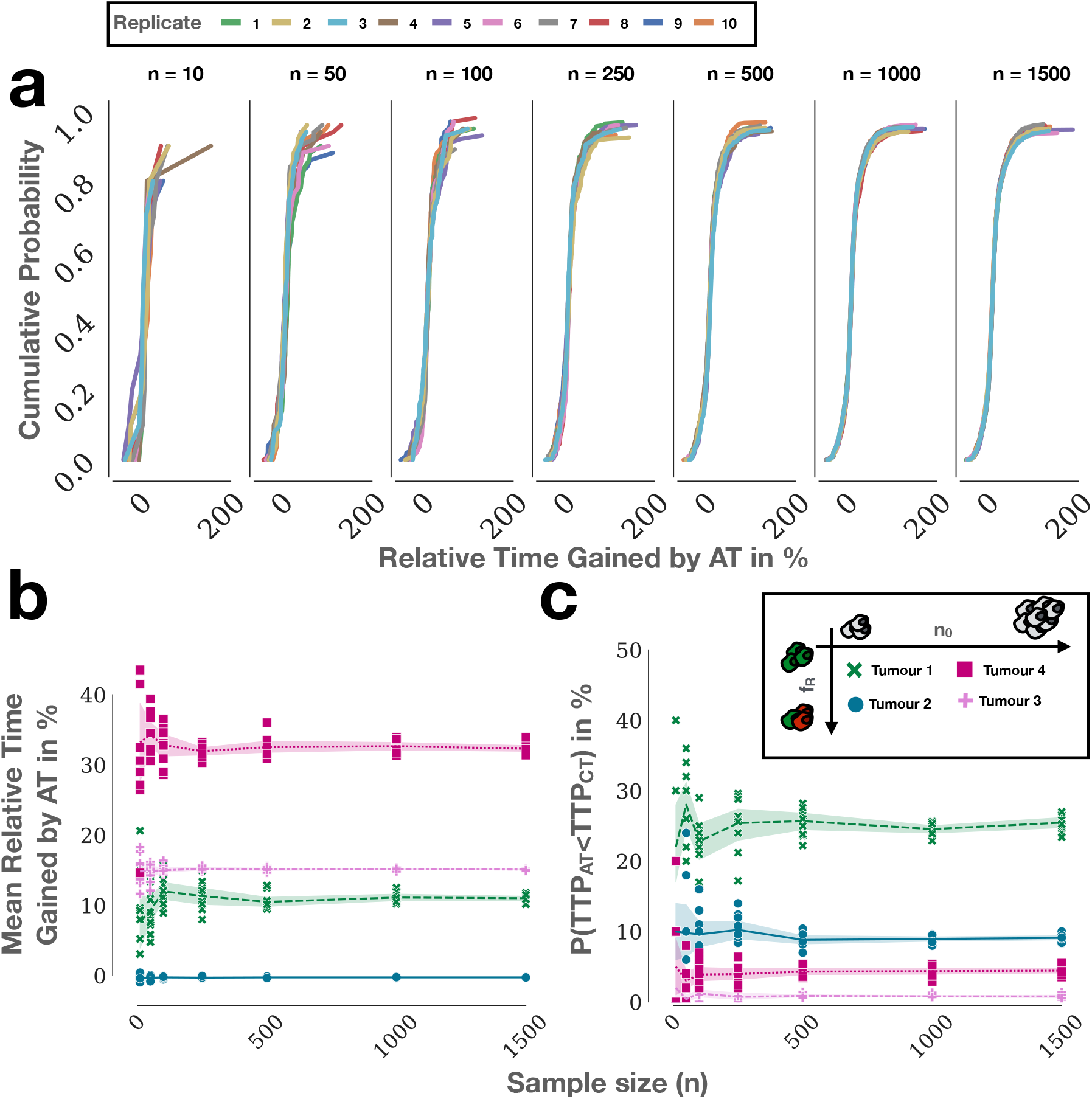
Representative results of the consistency analysis: (**a**) Increasing consistency in the distribution of time gained by adaptive therapy for one parameter set ((*n*_0_, *f_R_*, *c_R_*, *d_T_*) = (25%, 0.1%, 0%, 30%)) as the number of independent replicates per sample, n, is increased. For each value of n, 10 independent samples of sample size *n* were collected. (**b**) Mean value of the outcome distribution as a function of sample size, n, for four parameter sets (Tumour 1: (*n*_0_, *f_R_*) = (25%, 0.1%); Tumour 2: (*n*_0_, *f_R_*) = (25%, 10%); Tumour 3: (*n*_0_, *f_R_*) = (75%, 10%); Tumour 4: (*n*_0_, *f_R_*) = (75%,0.1%)). Lines indicate the mean value, shading a 95% confidence interval. Markers show values of individual replicates. For sample sizes upwards of 250, the different replicates produce very consistent results. (**c**) Proportion of runs in which adaptive therapy failed as a function of the sample size, *n*, for the same four parameter sets. Again for *n* ≥ 250 we see consistent values between independent replicates indicating that a sample size *n* ≥ 250 will yield a representative outcome distribution.

**Figure A2.**
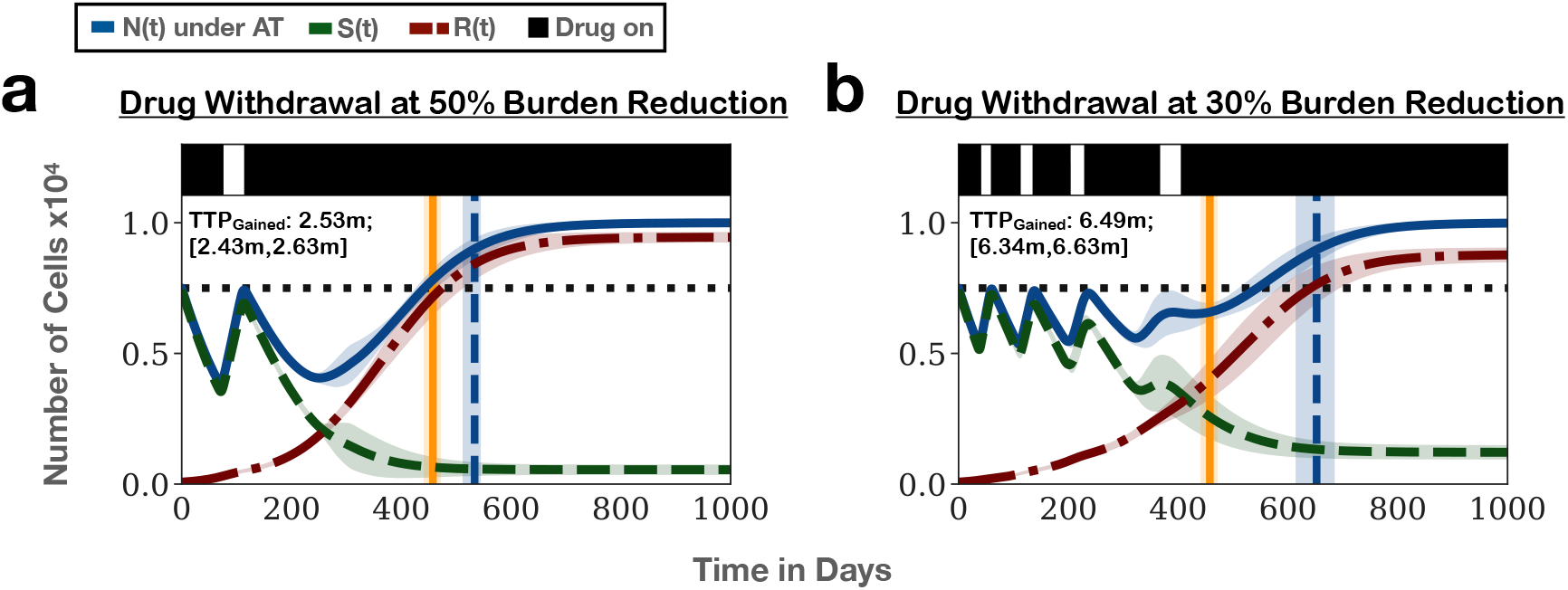
Comparison of two adaptive therapy protocols with different thresholds for treatment withdrawal. (**a**) Treatment withdrawal after a 50% tumour burden reduction. Lines and shading show the mean and standard deviation of the number of cells in each subpopulation and the total tumour size, respectively. Vertical lines denote the time of progression under continuous (yellow) and adaptive therapy (blue). Inset gives mean time gained and 95% confidence interval. (**b**) Treatment withdrawal after a 30% tumour burden reduction. This suggests that less aggressive treatment, especially if resistance is prevalent prior to treatment, will result in longer tumour control, and matches the results obtained for non-spatial models [15,16,22]. Note though that we are not taking into account possible risks associated with the higher tumour burden under the 30% threshold algorithm. Parameters: (*n*_0_,*f_R_*, *c_R_*, *d_T_*) = (75%, 1%, 0%, 0%); *n* = 250 independent replicates.

**Figure A3.**
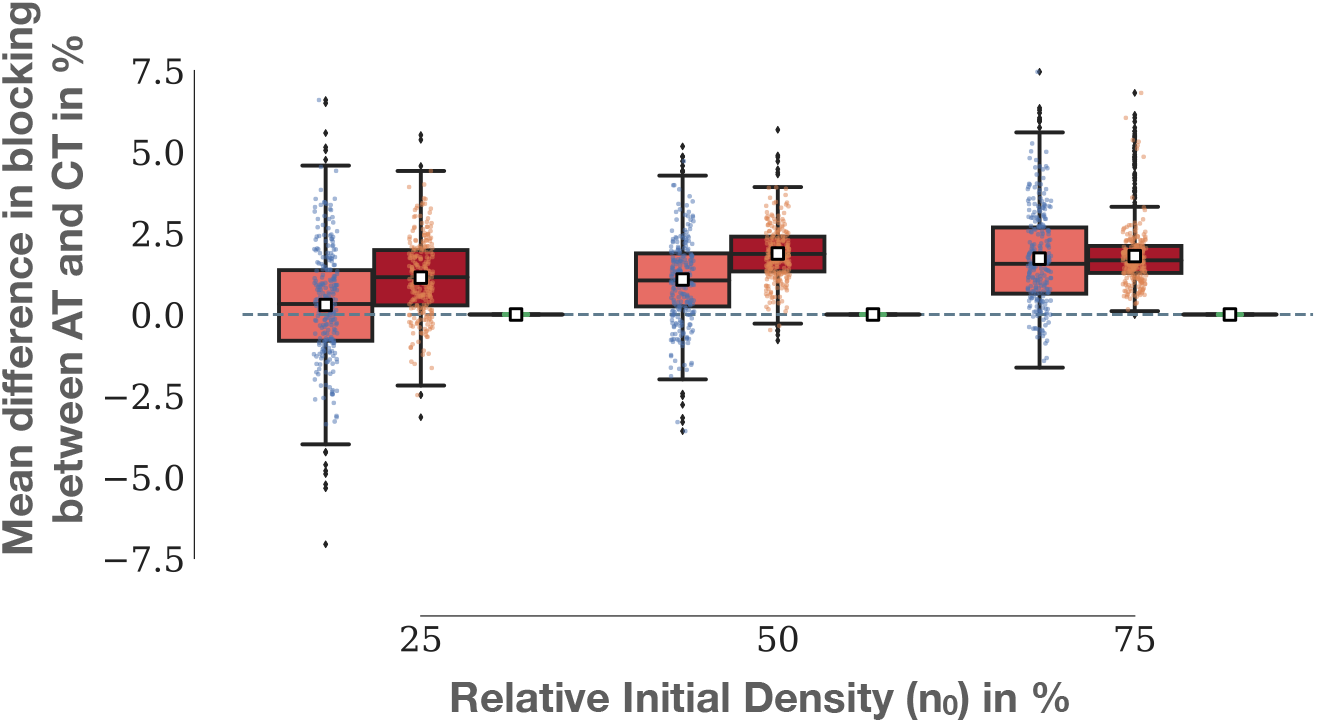
High density and low resistance fractions maximise the difference in the rates of competitive growth inhibition between continuous and adaptive protocols. Shown is the mean difference in the rate at which resistant cell divisions are blocked by continuous and adaptive therapy in the simulations in Figure 2e, given by 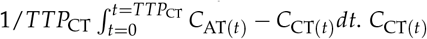 and *C*_AT(*t*)_ denote the rate of blocking under adaptive and continuous therapy as illustrated in Figure 2d. Parameters: (*n*_0_,*f_R_*, *c_R_*, *d_T_*) = (75%, 0.1%, 0%, 0%); 1000 independent replicates per condition.

**Figure A4.**
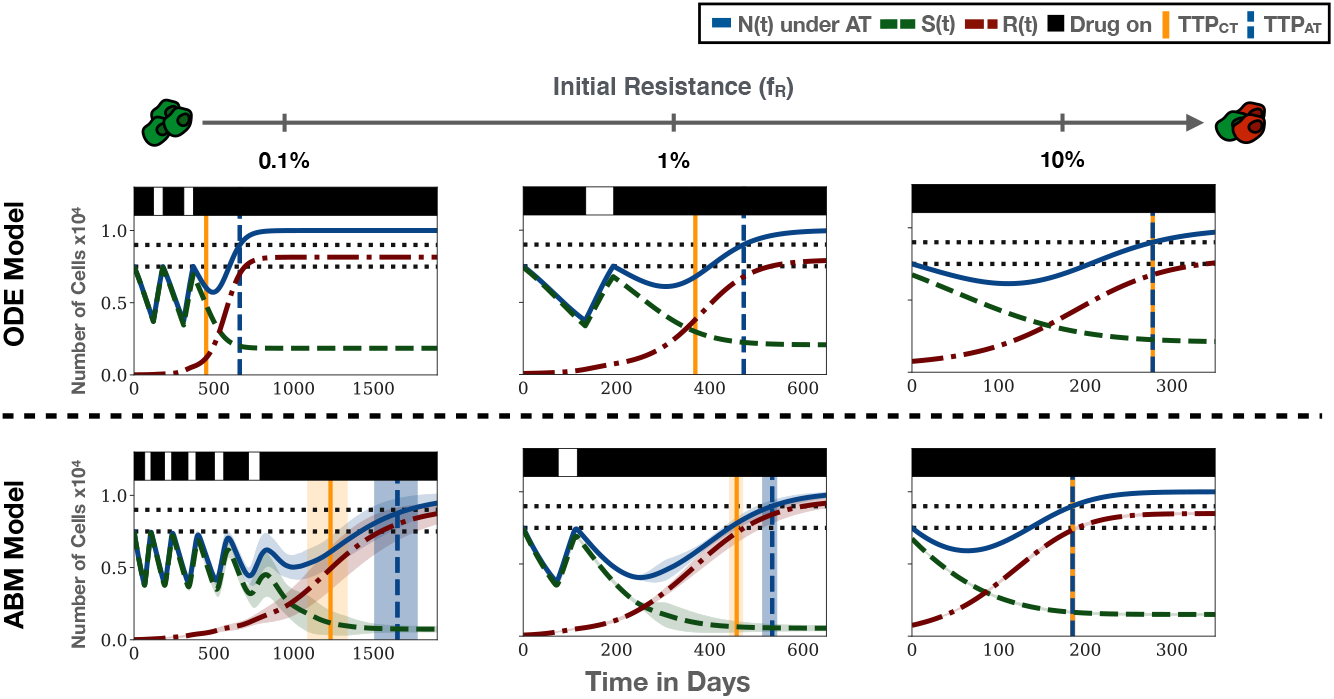
Comparison of the treatment predictions of the ABM and ODE model (Equations (2)-(4)) for the same initial density (*n*_0_ = 75%) but different initial resistance fractions. We assume no cost or turnover. For the ABM the mean and standard deviation of *n* = 250 independent replicates are shown. When the initial resistance fraction is small, the ABM predicts significantly longer TTP than the ODE model. When the resistance fraction is large, the converse is true. This indicates that because of the impact of space, different initial numbers of resistant cells (and thus independent colonies) result in distinct progression dynamics in the ABM.

**Figure A5.**
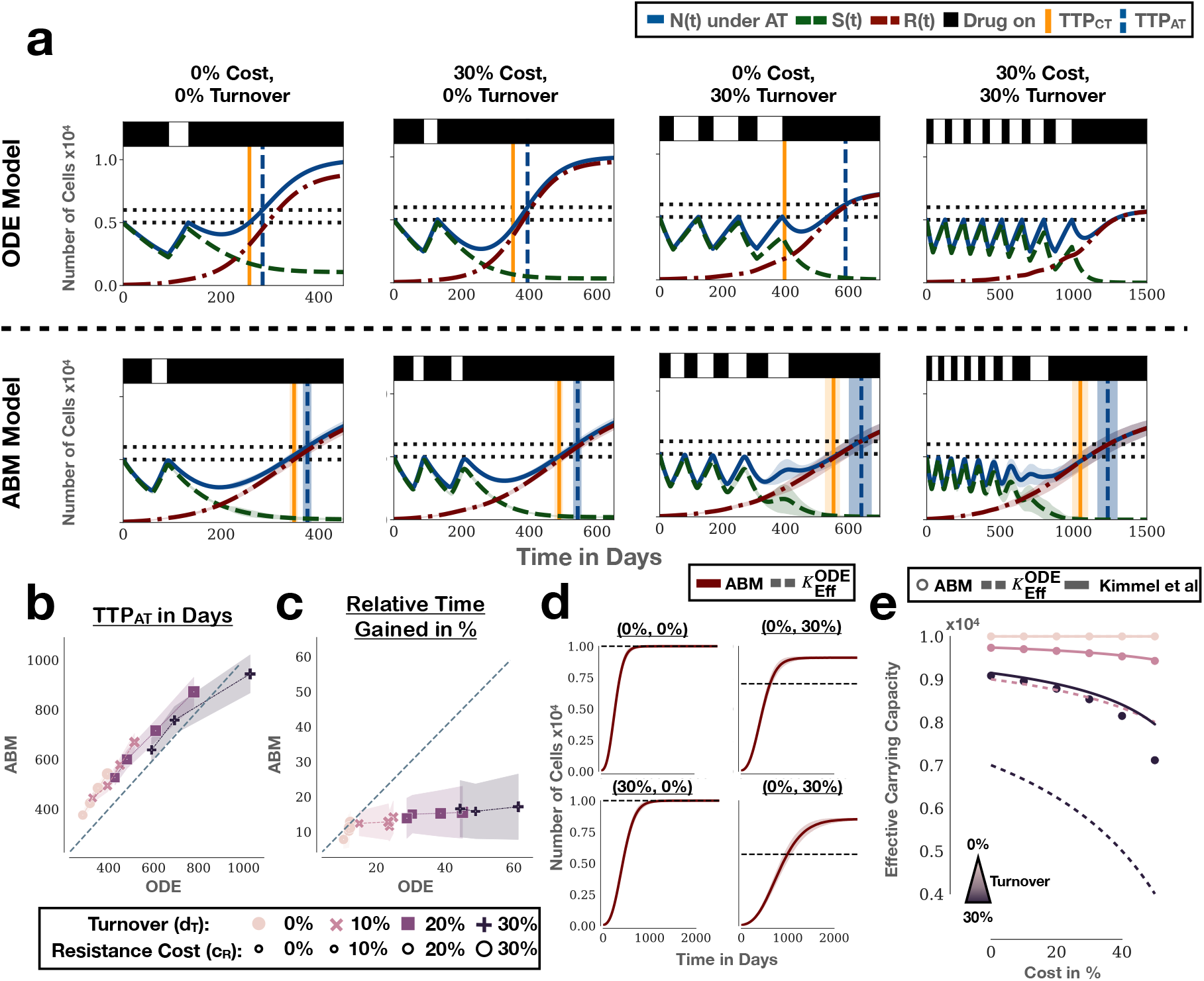
Comparison of the treatment predictions of the ABM and ODE model (Equations (2)-(4)) for different values of cost and turnover. (**a**) Matched simulations for different combinations of cost and turnover values ((*n*_0_, *f_R_*) = (50%, 1%)). For the ABM, the mean and standard deviation of *n* = 250 independent replicates are shown. (**b**) Comparison of the TTP under adaptive therapy in the ODE and ABM. The ABM predicts, in general, later progression but this difference becomes smaller in the presence of cost and turnover (*n* = 1000 replicates of the ABM). (**c**) Comparison of the relative time gained by adaptive therapy ((TTP_AT_-TTP_CT_)/TTP_CT_) predicted by the two models. The ABM predicts a significantly smaller benefit for adaptive therapy from cost and turnover than the ODE model. (**d**) The reason for the discrepancy observed in (c) is that the impact of turnover on population growth is smaller in the ABM than in the ODE. This can be seen by the fact that the effective carrying capacities, *K*_Eff_, (the steady states of the resistant population) are different between the two models in the presence of turnover. Shown are simulations of resistant cells grown in isolation (no drug) under different conditions (*n* = 250 replicates). The dashed line gives the steady state expected in the ODE model 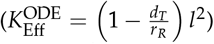. We previously showed that *K*_Eff_ is an important factor in determining the benefit of adaptive therapy [22]. As such, the differences in *K*_Eff_ explain the discrepancy in the predicted benefit of adaptive therapy. (**e**) The reason why the effective carrying capacities differ is the fact that in the ABM a cell has four potential neighbouring sites into which it can divide. This allows for more division to take place than predicted by the ODE model. To show this, we compare the effective carrying capacity in the ABM for different values of cost and turnover to an expression derived by Kimmel et al (in prep.) which accounts for the neighbourhood size: 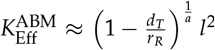, where *a* is the neighbourhood size (*a* = 4 for a von Neumann neighbourhood). With the neighbourhood size taken into account we see excellent agreement with the observed effective carrying capacity in the ABM. Values for the ABM were obtained by taking the final population size after 10y, as in (d) (*n* = 1000 replicates per condition).

**Figure A6.**
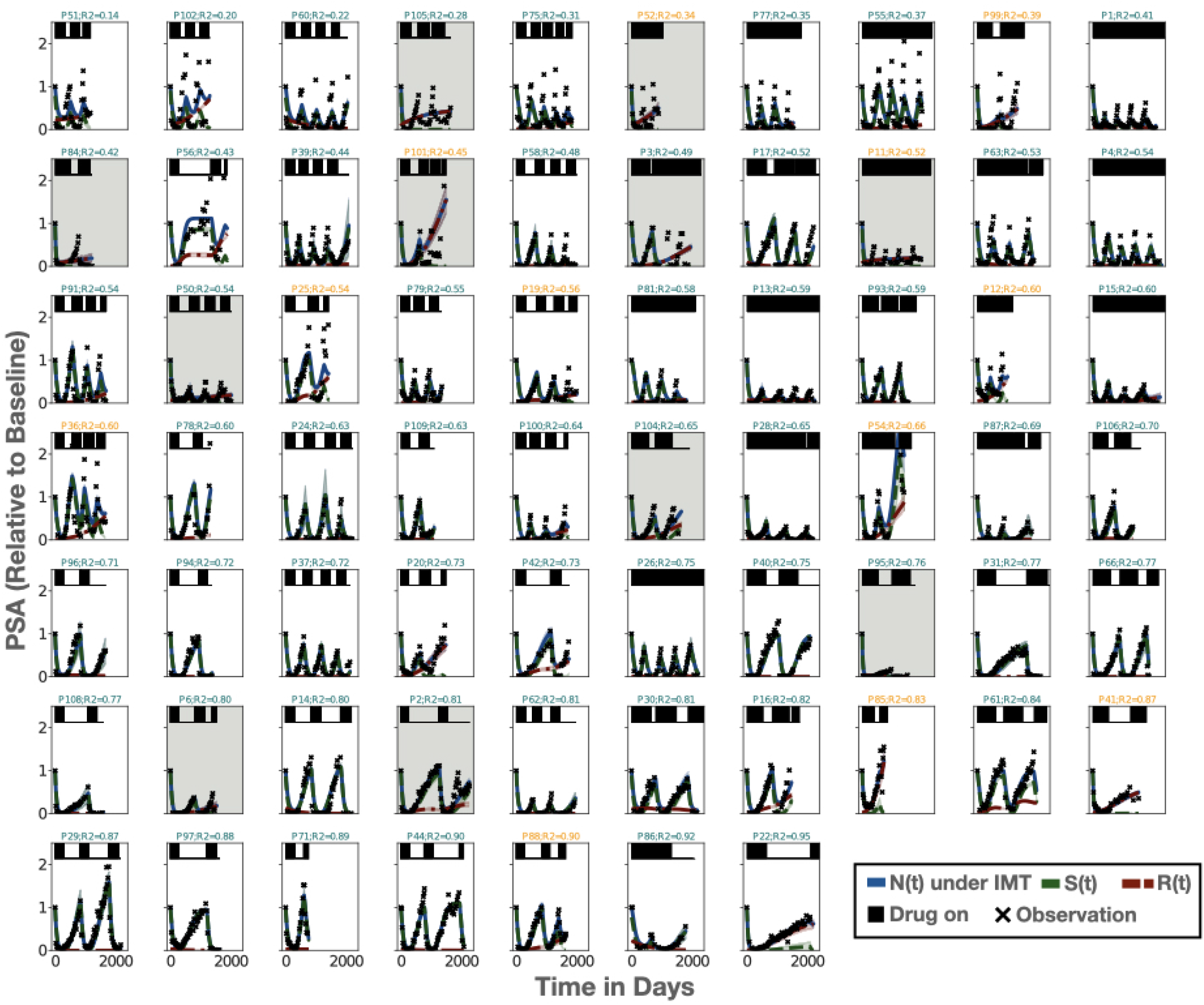
Overview of the ABM fits of the full model (all 4 parameters) for all 67 patients, arranged by their *r^2^* value (showing the mean and standard deviation of 25 replicates per patient). Title colour indicates whether a patient relapses (orange) or not (green). Patients who were excluded from further analysis due to poor model fits are marked with a grey background.

**Figure A7.**
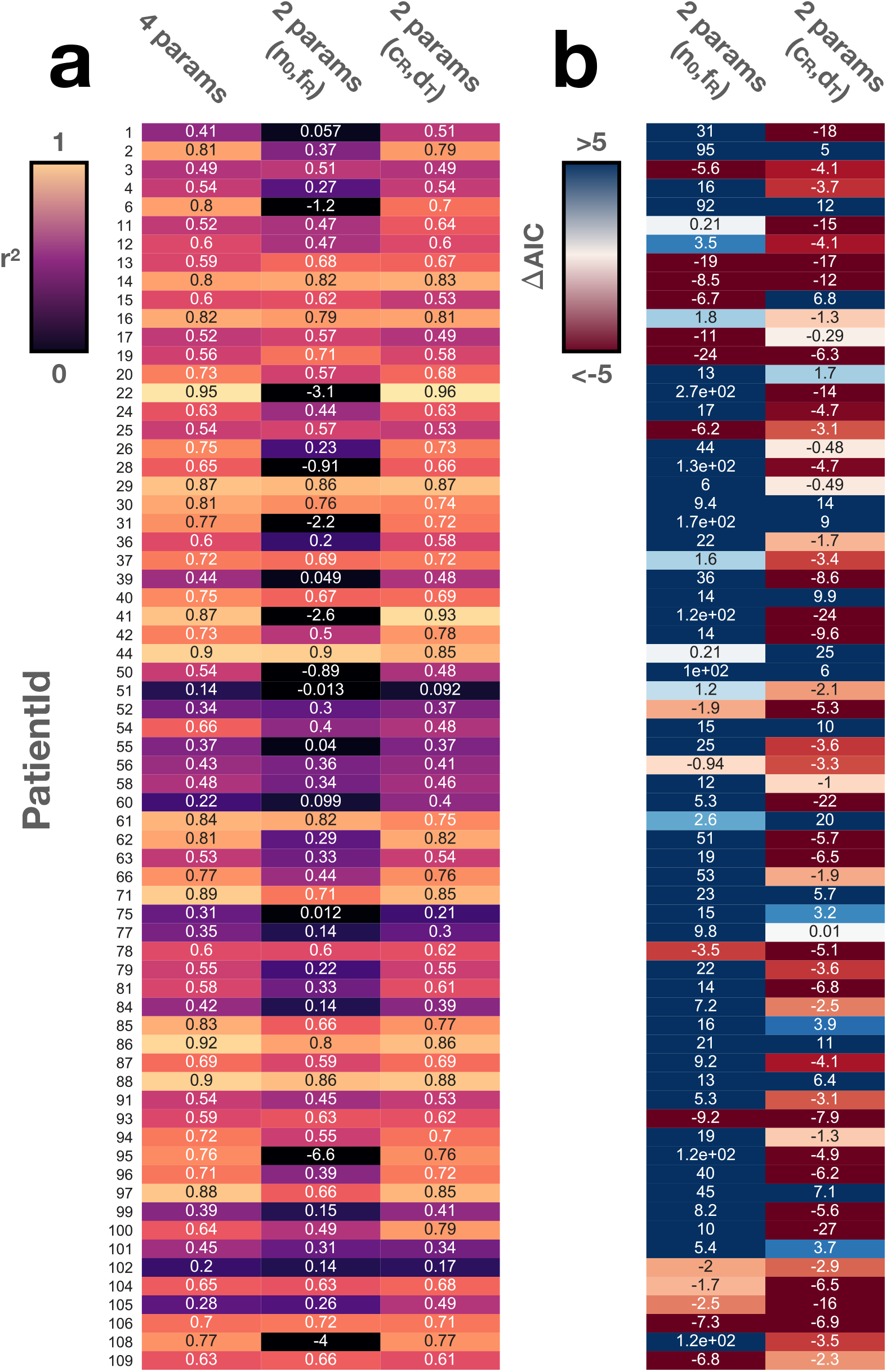
Comparison of fitting the ABM by allowing either the initial tumour composition (*n*_0_, *f_R_*), the cell kinetics (*c_R_, d_T_*), or all four parameters to vary (“4 params”). (**a**) Comparison of the *r^2^* values for each patient for each model. We observe that allowing only cost and turnover to be patient-specific can explain the data almost as well as the full 4 parameter model. This is not true for the model in which the initial conditions are patient-specific. (**b**) Difference in AIC between the 2 parameter models compared to the 4 parameter models for each patient. The AIC represents a measure of goodness-of-fit relative to a model’s complexity. When the AIC of two models differs by more than 2, the model with the smaller AIC is the preferred one [64]. This corroborates that for most patients the cost-turnover model provides the best description of the data.

**Figure A8.**
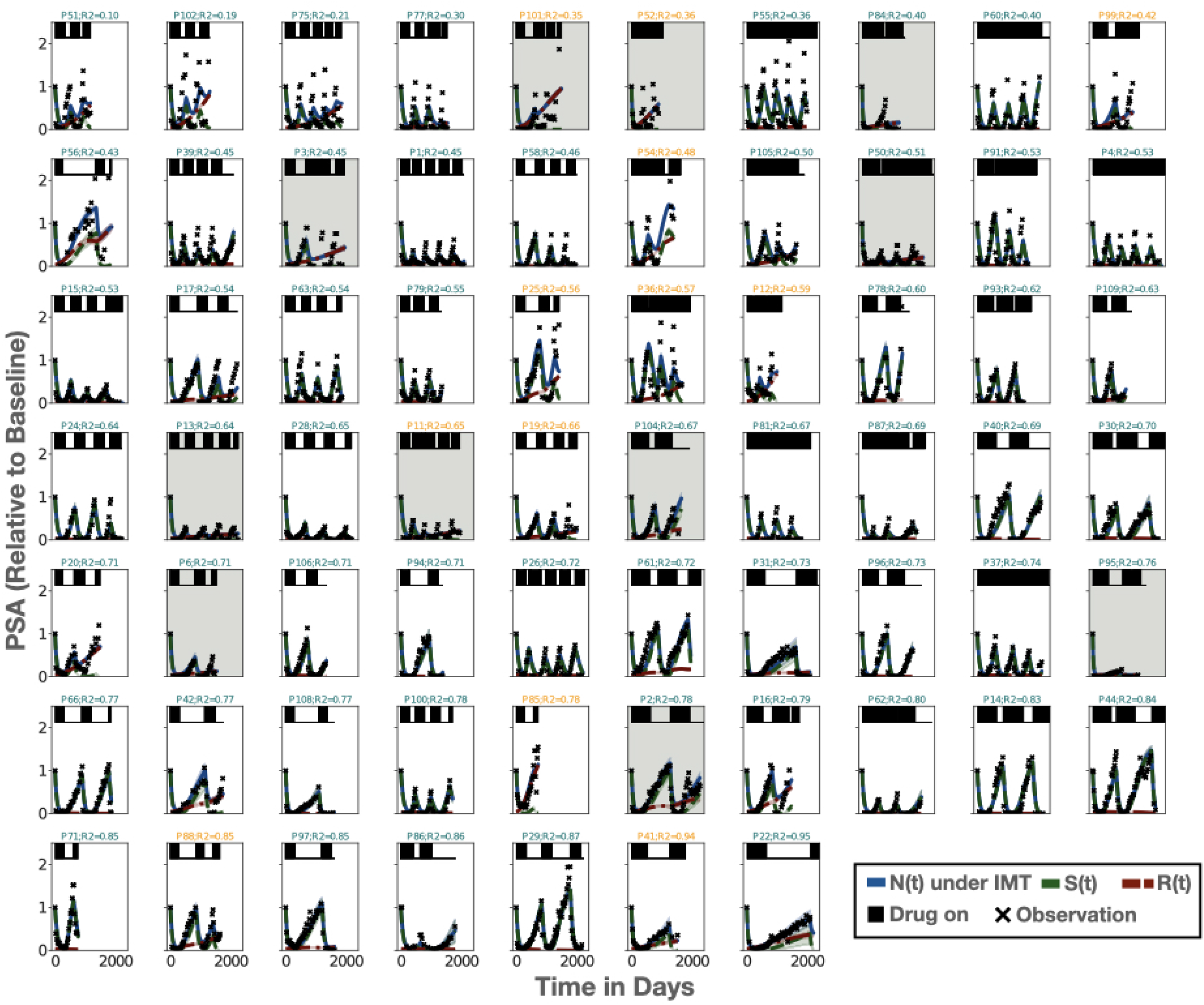
Overview of the ABM fits of the reduced model (fitting only cost and turnover) for all 67 patients, arranged by their *r^2^* value (showing the mean and standard deviation of 25 replicates per patient). Title colour indicates whether a patient relapses (orange) or not (green). Patients who were excluded from further analysis due to poor model fits are marked with a grey background.

